# Methods for decoding cortical gradients of functional connectivity

**DOI:** 10.1101/2023.08.01.551505

**Authors:** Julio A. Peraza, Taylor Salo, Michael C. Riedel, Katherine L. Bottenhorn, Jean-Baptiste Poline, Jérôme Dockès, James D. Kent, Jessica E. Bartley, Jessica S. Flannery, Lauren D. Hill-Bowen, Rosario Pintos Lobo, Ranjita Poudel, Kimberly L. Ray, Jennifer L. Robinson, Robert W. Laird, Matthew T. Sutherland, Alejandro de la Vega, Angela R. Laird

## Abstract

Macroscale gradients have emerged as a central principle for understanding functional brain organization. Previous studies have demonstrated that a principal gradient of connectivity in the human brain exists, with unimodal primary sensorimotor regions situated at one end and transmodal regions associated with the default mode network and representative of abstract functioning at the other. The functional significance and interpretation of macroscale gradients remains a central topic of discussion in the neuroimaging community, with some studies demonstrating that gradients may be described using meta-analytic functional decoding techniques. However, additional methodological development is necessary to fully leverage available meta-analytic methods and resources and quantitatively evaluate their relative performance. Here, we conducted a comprehensive series of analyses to investigate and improve the framework of data-driven, meta-analytic methods, thereby establishing a principled approach for gradient segmentation and functional decoding. We found that a two-segment solution determined by a k-means segmentation approach and an LDA-based meta-analysis combined with the NeuroQuery database was the optimal combination of methods for decoding functional connectivity gradients. Finally, we proposed a method for decoding additional components of the gradient decomposition. The current work aims to provide recommendations on best practices and flexible methods for gradient-based functional decoding of fMRI data.

## Introduction

The main goal of parcellating functional connectomes is to reduce high-dimensional connectivity space into a mosaic of brain regions. These regions condense the information from thousands of elements (i.e., voxels or vertices) that, on average, show coherent fluctuations in spontaneous activity during resting-state experiments (Bijsterbosch et al., 2020). Both soft-parcellation (e.g., independent component analysis: ICA) and hard-parcellation (e.g., boundary mapping and clustering) approaches have been established as popular methods for defining brain functional organization (Eickhoff et al., 2018). However, recent methodological developments for nonlinear dimensionality reduction techniques, using diffusion map embedding (Coifman et al., 2005) on functional magnetic resonance imaging (fMRI) matrices, have allowed the mapping of functional connectomes to low dimensional manifolds known as gradients of connectivity (Vos de Wael et al., 2020). This method aims to reduce the dimensionality of the data, similar to parcellation approaches; however, such reduction does not require discrete parcels, which generally remove the spatial transition between different brain regions (Huntenburg et al., 2018). Thus, the individual elements of the data (i.e., voxels or vertices) are preserved. Despite conceptual and mathematical differences between gradients and parcellations (Hong et al., 2020), it has been shown that the principal gradients of connectivity performed similarly to parcellations as behavioral predictors (Kong et al., 2023). Notably, gradients of functional connectivity complement parcellation methods by capturing a smoother transition between different functional regions, offering potential insight into hierarchical information processing in the brain (Margulies et al., 2016).

In a seminal publication, Margulies and colleagues demonstrated that the principal gradient of connectivity exists in the human brain, with unimodal primary sensorimotor regions situated at one end and transmodal regions associated with the default mode network and representative of abstract functioning at the other end (Margulies et al., 2016). That work provided a foundation for understanding macroscale gradients within the framework of functional hierarchies. Since then, gradient methods have been used to describe the continuous axis of functional specialization in regions such as the hippocampus (Vos de Wael et al., 2018) and insula (Royer et al., 2020; Tian and Zalesky, 2018). Gradient methods have also provided novel insight into disrupted connectivity among clinical populations (Hong et al., 2019), the organization of neonatal connectomes (Larivière et al., 2020a), and the description of functional connectivity changes after structural alterations (e.g., focal lesions) (Bayrak et al., 2019). However, the behavioral interpretation of macroscale gradients is still a central topic of discussion in the community.

Prior work has demonstrated that the functional significance of gradients may be described using meta-analytic functional decoding techniques (Margulies et al., 2016; Paquola et al., 2019). Using the Neurosynth database (Yarkoni et al., 2011), meta-analytic decoding of a connectivity gradient has previously been performed on connectivity gradient maps following two steps (Margulies et al., 2016). First, the gradient spectrum is segmented into five-percentile increments (Caciagli et al., 2022; Cross et al., 2021; Larivière et al., 2020b; Margulies et al., 2016; Murphy et al., 2018; Paquola et al., 2019), and the 20 resultant maps are binarized and transformed to volume space (Margulies et al., 2016; Paquola et al., 2019). Second, meta-analytic decoders, trained with term-based (Yarkoni et al., 2011) or Latent Dirichlet allocation (LDA) topic-based (Poldrack, 2011) meta-analyses from large-scale coordinate-based databases, are applied to each of the resultant maps. This decoding approach allows characterization of the full gradient spectrum, thereby addressing limitations of standard meta-analytic decoding, which would yield associations to only the vertices located at the higher end and leave the lower-end vertices unexplained. Although this approach has been used in multiple studies (Margulies et al., 2016; Paquola et al., 2019), additional methodological development is necessary to evaluate these procedures fully, including the segmentation approach and decoding strategy.

First, existing segmentation procedures are arbitrarily determined with equally dense intervals (i.e., five-percentile increments) (Caciagli et al., 2022; Cross et al., 2021; Larivière et al., 2020b; Margulies et al., 2016; Murphy et al., 2018; Paquola et al., 2019) and could be based on a more data-driven approach. In some cases, the segment’s boundary may fall in a high-density area (i.e., a point in the gradient space with many vertices in its vicinity), thus splitting a highly interconnected group of vertices into two different segments. As a result, vertices at the boundary of the gradient’s coordinate system may show a low confidence value. Although the vertices are close in gradient space to the vertices of their corresponding segment (i.e., highly interconnected), they are as close to the neighboring vertices from the boundary of the subsequent segment.

Second, the existing decoding approach transformed the gradient maps from the cortical surface to volume space, which required binarized maps, because surface-to-volume mapping is ill-defined for continuous maps since not every MNI voxel has a corresponding `fsaverage` vertex (Markello et al., 2022; Wu et al., 2018). Binarization removes the continuous representation of connectivity within each segment, eliminating the within-segment organization, which may remove important information and thus distort functional decoding results. Therefore, a meta-analytic decoding method implemented in the cortical surface, the native space of the gradient, should overcome this limitation.

Third, multiple different meta-analytic approaches for functional decoding are available as potential strategies for decoding connectivity gradients. Previously, gradient decoding has been conducted with term-based meta-analytic maps using 24 terms from Neurosynth, covering a limited range of cognitive functions (Paquola et al., 2019). Although such maps often approximate the results of manual meta-analyses of the same domain and produce accurate classifications of specific stimuli, that approach comes with redundant and ambiguous terms (de la Vega et al., 2016). Given the heterogeneous terminology across the literature, term-based meta-analyses from Neurosynth may map terms of the same construct (e.g., calculation, arithmetic, addition, and computation) to different maps (Dockès et al., 2020); thus, decoding based on term-based maps may yield association to only a few terms of a whole construct.

Alternatively, gradient decoding has used topic-based meta-analytic maps generated using a Latent Dirichlet allocation (LDA) model of publication abstracts in Neurosynth (Margulies et al., 2016). In this case, the “*v3-topics-50*” version of the database was used, including 50 topics extracted from an LDA of Neurosynth abstracts as of September 2014. Standard topic approaches provide a broad collection of meta-analytic maps associated with not just one but different terms (Poldrack et al., 2012), reducing the redundancy and ambiguity of term-based meta-analysis (de la Vega et al., 2016). In addition, the topic model is likely to group similar terms under the same construct and thus under the same meta-analytic map, thus overcoming the limitation of term-based meta-analysis with similar terms. However, the topics that a standard LDA produces are not constrained by neural data because the model works only on the text of publications (Rubin et al., 2017). In addition, the spatial components of the maps associated with each topic may be widely distributed and not reflect associations with clearly localized brain regions. Recently, a new decoding framework was introduced based on the Generalized Correspondence LDA (GC-LDA), an extension of the LDA model (Rubin et al., 2017). The GC-LDA model adds spatial and semantic constraints to the data, which allows the model to generate topic maps with a maximized correspondence between cognition and brain activity. However, such models have not yet been applied to decode functional connectivity gradients, and to date, no formal comparison across decoding strategies has been investigated.

The overall objective of the current study was to investigate and improve the framework of data-driven methods for decoding the principal gradient of functional connectivity, thereby promoting best practices for understanding its underlying mechanisms. We comprehensively examined and evaluated different methods to address the above limitations and establish a principled approach for gradient segmentation and meta-analytic decoding. To this end, we used the resting-state fMRI (rs-fMRI) group-average dense connectome from the Human Connectome Project (HCP) S1200 data release (Smith et al., 2013) to identify the principal gradient of functional connectivity. We evaluated three segmentation approaches: (i) percentile-based (PCT), (ii) segmentation based on a 1D *k*-means clustering approach (KMeans), and (iii) segmentation based on the Kernel Density Estimation (KDE) curve of the gradient axis. We assessed six different decoding strategies that used two meta-analytic databases (i.e., Neurosynth (Yarkoni et al., 2011) and NeuroQuery (Dockès et al., 2020)) and three methods to produce meta-analytic maps (i.e., term-based, LDA-based, and GC-LDA-based decoding). First, we evaluated the three segmentation approaches for 31 segmentation solutions using silhouette scores, variance ratio, and cluster separation. Then, we assessed the performance of the 18 meta-analytic segmentation/decoding strategies using multiple metrics, including correlation profile (CP), two semantic similarity measures, and signal-to-noise ratio (SNR). Finally, we performed a multidimensional decoding for the first four gradients.

We hypothesized that a data-driven segmentation approach (e.g., KMeans or KDE) is preferable to a segmentation based on an arbitrary percentile. In addition, we predicted that a continuous decoding approach based on a topic model (e.g., LDA or GC-LDA) would produce a broad set of meta-analytic maps with high association to each segment of the gradient, providing a reliable functional description and aiding overall interpretability. Further, we expected that the terms provided by such topic-based decoders would yield a high information content and SNR compared to other strategies. The current work recommends best practices and flexible methods for gradient-based functional decoding of fMRI data.

## Materials and Methods

Our analysis plan was preregistered at https://osf.io/5q29z. We conducted additional analyses during the review process that deviated from the original protocol, as suggested by the reviewers. These deviations supplemented and enhanced the original analysis plan but did not substantially alter our initial aims and hypotheses. **Figure 1** provides an overview of our methodological approach. First, we performed gradient decomposition of the group-average dense connectome from HCP resting-state fMRI data. Second, we evaluated three different segmentation approaches to split the gradient spectrum into a finite number of brain maps. Third, we implemented six different decoding strategies, for a total of 18 decoding strategies, when combined with each segmentation approach. Fourth, we evaluated the decoding strategies using multiple metrics to compare relative performance. Fifth, and as a final step, we proposed a method for decoding additional components of the gradient decomposition. Subsequent sections describe these five steps in more detail.

**Figure 1.**
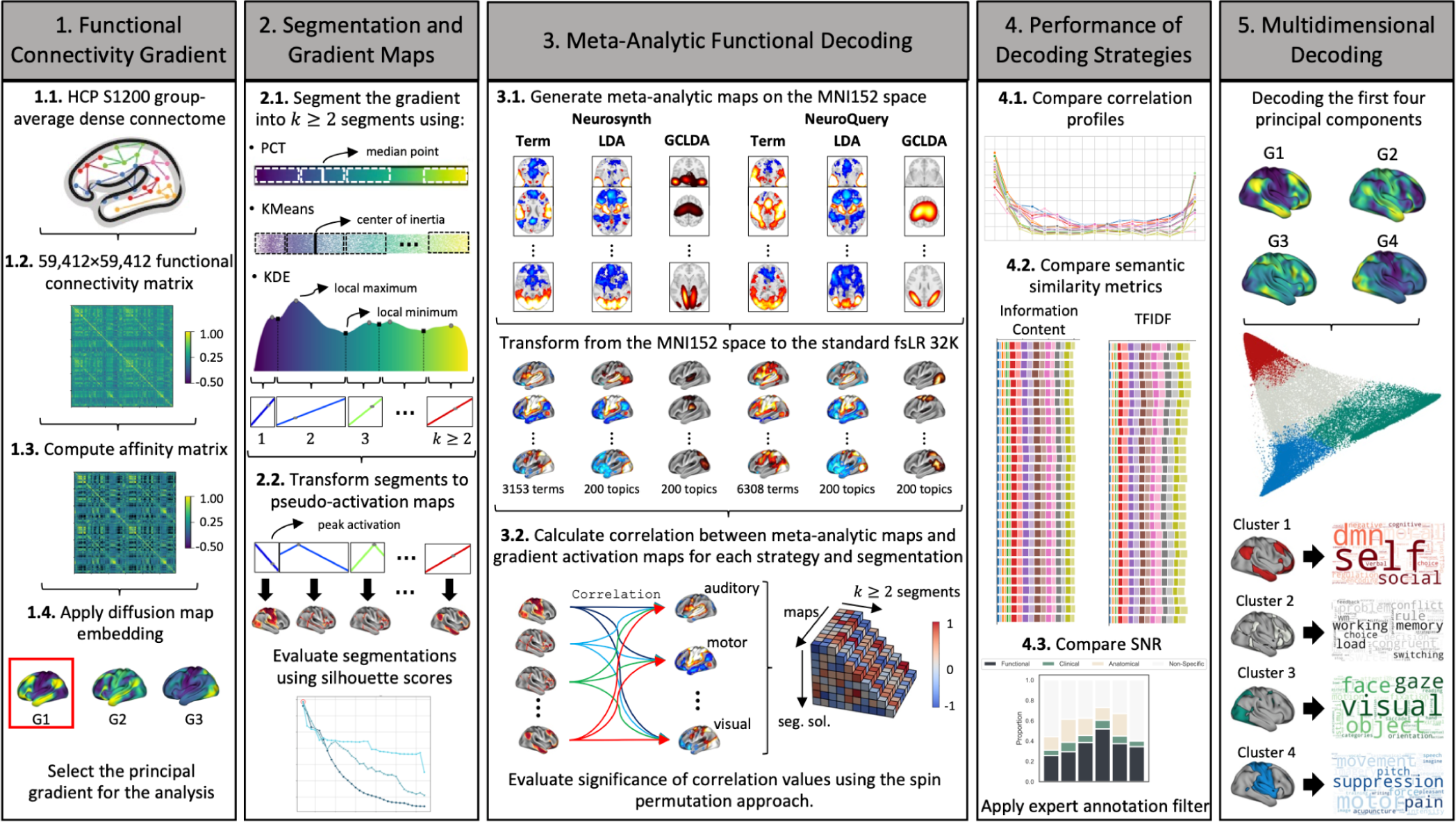
Workflow for Evaluating Meta-Analytic Functional Decoding of the Functional Connectivity Gradient. HCP S1200 resting-state fMRI data were used (**Step 1.1**) to generate functional connectivity matrices (**Step 1.2**) and compute the affinity matrix (**Step 1.3**). Diffusion map embedding was applied to identify the principal gradient of functional connectivity (**Step 1.4**). Whole-brain gradient maps were segmented to split the gradient spectrum into a finite number of brain maps. Three different segmentation approaches were evaluated: percentile-based (PCT), *k*-means (KMeans), and KDE (**Step 2.1**). Individual segments were transformed into “activation” brain maps for decoding (**Step 2.2**). The three segmentation approaches were evaluated using the silhouette scores. Six different meta-analytic decoding strategies were implemented in surface space, derived from three sets of meta-analytic maps (i.e., term-based (Term), LDA, and GC-LDA) and two databases (i.e., Neurosynth and NeuroQuery) (**Step 3.1**). Results from the decoding strategies were evaluated using four performance metrics, assessed by comparing correlation profiles (**Step 4.1**), semantic similarity metrics (i.e., information content and TFIDF) (**Step 4.2**), and signal-to-noise ratio (SNR) (**Step 4.3**). In addition, we remove the non-functional terms from the model. Finally, we performed a multidimensional decoding using the first four components together **(Step 5**).

### Analysis Step 1: Functional Connectivity Gradient

#### Connectivity Data

We used the rs-fMRI group-average dense connectome from the Human Connectome Project (HCP) S1200 data release (Smith et al., 2013; Uğurbil et al., 2013; Van Essen et al., 2013, 2012). The rs-fMRI experiment consisted of four runs of approximately 14.5 minutes (1,200 frames) each, for a total of 4,800 frames of resting-state data per participant. 1,003 subjects that completed all four rfMRI runs (i.e., their resting-state data comprising 4,800 frames) were included in the analysis. Each run was preprocessed using HCP pipelines (Fischl, 2012; Glasser et al., 2013; Jenkinson et al., 2012, 2002) and denoised using ICA-FIX (Feinberg et al., 2010; Glasser et al., 2016; Salimi-Khorshidi et al., 2014). The data were temporally demeaned, and its variance was normalized according to (Beckmann and Smith, 2004). The cerebral cortex between subjects was coregistered using the Multimodal Surface Matching algorithm (MSMAll). Then, group-PCA from the 1,003 subjects, performed by MELODIC’s Incremental Group-PCA (MIGP), produced 4,500 weighted spatial eigenvectors. These eigenvectors were renormalized, eigenvalue-reweighted, and correlated to produce the dense connectome (59, 412×59, 412 z-transformed functional connectivity matrix) (**Fig. 1, Step 1.1**). Further details on this approach can be found at www.humanconnectome.org/storage/app/media/documentation/s1200/ HCP1200-DenseConnectome+PTN+Appendix-July2017.pdf. Following Margulies et al. (Margulies et al., 2016), an inverse Fisher transform was applied to the z-transformed correlation to scale the values between -1 and 1 (**Fig. 1, Step 1.2**).

#### Gradient Decomposition

Diffusion embedding (Coifman et al., 2005) was applied to the group-averaged connectivity matrix using the `mapalign` Python package (Langs et al., 2015), capturing spatial gradients in macroscale resting-state functional organization. The dimensionality reduction algorithm was run using diffusion maps embedding. The procedure is detailed in Vos de Wael et al., 2020. It includes the following steps: (i) the input matrix was made sparse by retaining the top 10% of weighted connections per row, and a cosine similarity matrix was computed to capture similarity in connectivity matrices between vertices (**Fig. 1, Step 1.3**); (ii) diffusion map embedding was applied to identify principal gradient components explaining connectome variance (n_components x n_vertices: 9 x 59,412); and (iii) the amount of variance explained by each gradient was calculated (**Fig. 1, Step 1.4**). We used the gradient (i.e., eigenvector) with the highest variance explained (i.e., eigenvalue), also called the principal gradient, for the first part of the analysis. Additional lower-order cortical gradients were used to perform multidimensional decoding in the last analysis step. In the first analysis steps, we focus only on the principal gradient because it represents an indicator of the global connectivity of the graph (Lioi et al., 2021) and has been observed to be highly consistent across resting-state dimensionality decomposition studies (Caciagli et al., 2022; Hong et al., 2019; Margulies et al., 2016; Paquola et al., 2019).

### Analysis Step 2: Segmentation and Gradient Maps

Once diffusion embedding was applied to decompose the connectivity matrix into whole-brain gradient connectivity maps, the new coordinate system was examined by segmenting the gradient into individual maps in preparation for meta-analytic decoding. Here, we used “segment” for one-dimensional gradients and “cluster” for high-dimensional gradients.

Importantly, separating the gradients in sparse maps removes the main advantage of the gradient dimensionality reduction over parcellation approaches, wherein individual elements are preserved instead of grouping them in discrete parcels, thus capturing a smoother transition between different brain regions. However, in the context of meta-analytic decoding, analyzing sparse maps within the gradient axis is essential. Naturally, the most effective solution for decoding the principal gradient will be to characterize each element (i.e., vertex) separately or describe the axis as a single map. However, it is still impractical to decode brain maps at such extreme scales, given existing limitations of decoding algorithms, which are generally trained with sparse maps drawn from the neuroimaging literature (e.g., meta-analytic maps). First, finding a label associated with single vertices of the cortical surface is unfeasible. A single vertex can be activated in almost all meta-analytic maps from the training sample, preventing us from finding the most likely labels associated with the target vertex. Second, considering the dense map as a single target in the existing decoding algorithm will only produce associations with the vertices at one end of the gradient axis, leaving the other vertices unexplained. The values in a gradient represent coordinates in connectivity space. Regions with similar values (e.g., 4.1 and 4.2) are closer in space and thus more strongly connected, while regions that are further apart (e.g., -3.1 and 4.2) are weakly connected (Kong et al., 2019).

#### Segmentation of the Cortical Gradient

We evaluated three segmentation approaches: (i) percentile-based segmentation (PCT), (ii) segmentation based on a 1D k-means clustering approach (KMeans), and (iii) segmentation based on the Kernel Density Estimation (KDE) curve of the gradient axis (**Fig. 1, Step 2.1**). For percentile-based segmentation, we applied the previously implemented method (Margulies et al., 2016) in which the whole-brain gradient is segmented into percentile increments. Then, we applied two additional, data-driven segmentation approaches to minimize the effect of hard, arbitrary boundaries. The second segmentation approach relied on 1D k-means clustering, previously used to define clusters of functional connectivity matrices to establish a brain-wide parcellation (Yeo et al., 2011). The third segmentation approach was inspired by methods of cytoarchitectonic border detection using the gray level index (Bludau et al., 2014). This method identifies local minima of a Kernel Density Estimation (KDE) curve of the gradient axis, which are points with minimal numbers of vertices in their vicinity. The number of segments for the KDE method was modified by tuning the width parameter to generate the KDE curve.

To compare the three segmentation approaches (i.e., PCT, KMeans, and KDE), we estimated silhouette measures with the Python library `scikit-learn` (Abraham et al., 2014); the silhouette measure is useful for determining the confidence of each spatial location in cortical parcellations (i.e., “clusters”) (Yeo et al., 2011). Silhouette scores were used to generate confidence maps, which allow whole-brain visualization across different segmentation approaches. Different segmentations of the principal gradient were generated for each segmentation approach, corresponding to numbers of segments ranging from *k* = 2 to *k* = 32 for a total *n* = 31. Further, to determine average relative performance across segmentation approaches, we used three metrics that have been previously used for evaluating clustering performance in connectivity matrices (Bzdok et al., 2015; Eickhoff et al., 2016a, 2016b; Flannery et al., 2020; Morawetz et al., 2020). First, we determined the mean silhouette coefficient, computed by taking the average of the silhouette scores over all samples (Rousseeuw, 1987), the variance ratio (Calinski and Harabasz) score (Caliński and Harabasz, 1974), and the cluster separation (Davies-Bouldin) score (Davies and Bouldin, 1979). Definitions for these metrics can be found in `scikit-learn`. The highest mean silhouette score, highest variance ratio, and lowest cluster separation represent the best relative performance.

#### From Gradient Space to Pseudo-Activation Maps

Following segmentation, individual segments were transformed into pseudo-activation maps before meta-analytic decoding. In the original approach (Margulies et al., 2016), the gradient maps were binarized, removing the continuous representation of connectivity within each segment. Although the segments are still hierarchically organized, binarization eliminates the within-segment organization, which may remove important information and thus distort functional decoding results. We omitted the binarization step for the current study and leveraged the within-segment continuous axis to generate pseudo-activation maps. We remind readers that the values of the gradient maps represent coordinates in the low-dimensional manifold. Thus, to decode maps as if they were activation maps of continuous values, we computed the affinity of each element to a reference point with an inverse function of the distance as defined in the same gradient space.

First, we defined a peak activation point (or reference point) for each segmentation approach to minimize the effect of boundaries with continuous clusters. For the two segments located at the ends, the peak activation was defined on the farther boundary from the contiguous segment. This criterion only applies to the 1D case, where the terminal segments of the principal gradient are clearly defined in space. Meanwhile, the peak activation for intermediate segments was defined as follows. For PCT, we chose the median point within subsegments, while for KMeans, we selected the center of inertia as provided by the k-means algorithm. In addition, we thought that the local maximum better represented the peak activation for KDE. The extreme value theorem guarantees the existence of a maximum between two minima for a continuous function in a closed interval. Next, the pseudo-activation values were determined by affinity values (*A*(*v*, *p*)) from all elements (*v*) to the peak activation (*p*) location within each segment, using a Gaussian (radial basis function: RBF) kernel.

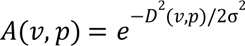

The distances (*D*(*v*, *p*)) were calculated in the gradient space using the Euclidean norm. The sigma coefficient (σ) was defined as the average distance within the segment. The final values range between 0 and 1, where 1 represents values closer in space or strongly connected to the reference point, and 0 indicates elements closer to the boundaries or weakly connected to the reference point. **Fig. 1, Step 2.2** illustrates how the process looks for 1D space (only including the principal gradient). The same transformation is generalizable to higher dimensional clustering with k-means when other gradients (e.g., the second and third gradient) of connectivity are included in the segmentation/clustering procedure.

It is worth noting that utilizing continuous maps on surface space required the functional decoding approach to be performed in surface space. Margulies et al.’s approach did not warrant consideration of this issue because it made use of binary maps in place of continuous maps (Margulies et al., 2016), which may be accurately transformed from surface to volume space using the nearest vertex approach (Wu et al., 2018). However, surface-to-volume mapping is ill-defined for continuous maps because not every MNI voxel has a corresponding `fsaverage` vertex (Markello et al., 2022; Wu et al., 2018).

### Analysis Step 3: Meta-Analytic Functional Decoding

Meta-analytic functional decoding provides a quantitative estimation of behavioral profiles associated with brain regions or networks (Amft et al., 2015; Bzdok et al., 2013b, 2013a; Cieslik et al., 2013; Laird et al., 2009; Nickl-Jockschat et al., 2015; Poldrack, 2011; Rottschy et al., 2013; Smith et al., 2009). This decoders are trained with a collections of maps generated by large-scale meta-analytic databases (e.g., Neurosynth (Yarkoni et al., 2011) and NeuroQuery (Dockès et al., 2020)). Margulies and others pioneered meta-analytic functional decoding of the connectivity gradient in volume space using the Neurosynth database (Margulies et al., 2016). However, multiple different meta-analytic decoding approaches are available as potential strategies for decoding gradients. In the current study, we assessed six different decoding strategies that used two meta-analytic databases (i.e., Neurosynth and NeuroQuery) and three sets of meta-analytic maps (i.e., term-based, LDA-based, and GC-LDA-based decoding) (**Fig. 1, Step 3.1**).

#### Meta-Analytic Databases

In neuroimaging, meta-analytic databases can be found in three different formats: (1) coordinate-based databases with automatically annotated studies, such as Neurosynth (Yarkoni et al., 2011) and NeuroQuery (Dockès et al., 2020), (2) coordinate-based databases with manually annotated studies, such as BrainMap (Laird et al., 2011b, 2011a, 2005), and (3) image-based databases such as NeuroVault (Gorgolewski et al., 2015).

Considering the trade-off between breadth and depth in the goal to develop a sufficiently large meta-analytic database that covers a broad range of mental functions (Varoquaux et al., 2018), in this work, we used large coordinate-based databases whose studies were automatically annotated, and provide a large set of cognitive states/tasks and activation coordinates. We used the popular Neurosynth database (Yarkoni et al., 2011), which contains activation coordinates automatically extracted from 14,371 published fMRI studies and associated 3,228 semantic terms from the corresponding article abstracts. In addition, we used the recently released NeuroQuery database (Dockès et al., 2020), which contains activation coordinates that were automatically extracted from 13,459 published fMRI studies, as well as associated 6,308 terms or phrases related to neuroscience extracted from the articles’ abstract, body, author-provided keywords, and title. We downloaded the database and features from the Neurosynth version 7 data release (github.com/neurosynth/neurosynth-data) and the NeuroQuery version 1 data release (github.com/neuroquery/neuroquery_data) using the `fetch_neurosynth` and `fetch_neuroquery` functions, respectively, from the NiMARE (Neuroimaging Meta-Analysis Research Environment) Python package (Salo et al., 2022a, 2022b). In both databases, each document text was represented with the term-frequency inverse document frequency (TFIDF) for each term of the vocabulary in the entire corpus of fMRI studies. The databases were converted to a NiMARE object using `convert_neurosynth_to_dataset` (Salo et al., 2022a). Then, we downloaded article abstracts from PubMed using the PubMed IDs with NiMARE’s function `download_abstracts` to create a corpus to train topic-based models.

#### Meta-Analytic Maps

Next, six sets of meta-analytic maps with associated terms were generated from the Neurosynth and NeuroQuery datasets. Meta-analytic maps are generated from these databases using coordinate-based meta-analysis (CBMA). Resultant meta-analytic maps are associated with a vocabulary of concepts related to cognitive states or tasks, which are then used for functional decoding. Each set of maps was generated using three different meta-analytic methods: term-based, LDA-based, and GC-LDA-based approaches. Term-based (Paquola et al., 2019) and LDA-based maps (Margulies et al., 2016) have previously been used for gradient decoding, while the GC-LDA method (Rubin et al., 2017) has not yet been applied to decode connectivity gradients.

#### Term-Based Meta-Analytic Maps

Term-based meta-analysis (Yarkoni et al., 2011) uses database-specific vocabulary terms, their frequency of occurrence across articles, and the activation coordinates of these articles in a dataset to create meta-analytic maps. As mentioned above, each term in the database is represented by the TFIDF across articles, which is used to identify studies where the term was frequently used. To create the meta-analytic sample of studies for a term, we applied a frequency threshold of 0.001 to eliminate studies that used the term incidentally. Study-specific maps were generated by convolving the peak activation with a binary sphere of 10mm radius centered at the voxel of each activation coordinate, then combining the map from each activation coordinate by taking the maximum value for each voxel. Next, meta-analytic maps were generated by fitting the meta estimator `MKDAChi2` class from NiMARE’s CBMA module to the resulting modeled activation maps from the meta-analytic sample. The chi-square estimator differs from the typical multilevel kernel density analysis (MKDA) (Kober et al., 2008; Wager et al., 2009, 2007) as it uses voxel-wise chi-squared tests to generate statistical maps showing the dependency between term and activation. Finally, for each database, we generated one meta-analytic map for each term from the vocabulary of each database; that is, 3,228 meta-analytic maps from Neurosynth and 6,145 meta-analytic maps from NeuroQuery.

#### LDA-Based Meta-Analytic Maps

The topic modeling technique, LDA, has been applied to text from published fMRI articles to identify latent structure and produce semantically coherent neuroimaging topics (Poldrack et al., 2012), revealing patterns consistent with previous meta-analyses (Poldrack et al., 2012; Rubin et al., 2017). LDA-based meta-analytic maps have been used for the functional decoding of brain networks (de la Vega et al., 2016; Wang et al., 2020) and individual segments of functional connectivity gradients (Margulies et al., 2016) as an alternative to overcome drawbacks associated with term-based meta-analytic methods.

In the current study, we performed topic-based meta-analyses using an LDA model on article abstracts from Neurosynth and article abstracts, bodies, author-provided keywords, and titles from NeuroQuery. We generated 200 different topics with two sets of probability distributions: (i) the probability of a word given topic *p(word|topic)* and (ii) the probability of a topic given article *p(topic|article)*. To generate topic-wise meta-analytic maps, we conducted a meta-analysis for each topic, where spatial mappings were indirectly computed via the documents’ topic loadings. Similar to term-based meta-analysis, we applied a frequency threshold of 0.05 to the probability *p(topic|article)* to eliminate articles that used the topic incidentally and created the meta-analytic sample of studies for a topic. Then, meta-analytic maps were generated by fitting the meta estimator `MKDAChi2` class from NiMARE’s CBMA module to each article’s resulting modeled activation maps of each article from the topic. Finally, for each database, we generated one meta-analytic map for each topic with the collection of words given by the top probability values of *p(word|topic)*; that is, 200 meta-analytic maps from Neurosynth and 200 meta-analytic maps from NeuroQuery.

#### GC-LDA-Based Meta-Analytic Maps

Recently, an extension to the LDA model, the GC-LDA, was introduced by Rubin et al. (Rubin et al., 2017). The GC-LDA model adds spatial and semantic constraints to the data, which allows the model to generate topic maps with a maximized correspondence between cognition and brain activity. This method, in addition to the two probability distributions from LDA (i.e., *p(word|topic)* and *p(topic|article)*), produces an additional probability distribution: (iii) the probability of each topic given a voxel: *p(topic|voxel)*. The default implementation of the GC-LDA decoding in NiMARE makes use of a dot product continuous decoding approach: *p(world|image)* = τ ·*p(world|topic)*, where *p(world|image)* is the vector of term/word weight associated with an input image *I* (e.g., unthresholded and standardized segment map) and τ = *p(topic|voxel)* · *I*(*voxel*) is the topic weight vector, *I*(*voxel*) is a vector with *z*-score value for each masked voxel of the input image. The term *p(world|image)* gives the most likely word from the top associated topics for a given unthresholded statistical map of interest. However, to keep the GC-LDA decoding strategy comparable to the other two strategies in terms of producing meta-analytic maps with high correlation across a segment of the gradient axis, we used the topic-wise meta-analytic maps given by the *p(topic|voxel)* distribution for functional decoding. Thus, similar to the LDA approach, the final step of the GC-LDA approach was to generate, for each database, one meta-analytic map for each topic with the collection of words given by the top probability values of *p(topic|voxel)*; that is, 200 meta-analytic maps from Neurosynth and 200 meta-analytic maps from NeuroQuery.

#### Continuous Decoding Approach

Ultimately, these procedures yielded six different decoding strategies derived from three sets of meta-analytic maps (i.e., term-based, LDA-based, GC-LDA-based) and two databases (i.e., Neurosynth and NeuroQuery). We then applied a continuous decoding approach to decode the segmented gradient maps from the three segmentation approaches. Continuous decoding targeted unthresholded statistical maps (e.g., unthresholded pseudo-activation maps from the gradient axis). It can be accomplished by either generating correlation coefficients between unthresholded meta-analytic maps and unthresholded statistical maps or computing the dot product between them. We used the first approach, which is implemented in NiMARE’s `CorrelationDecoder` class object (**Fig. 1, Step 3.2**). **Table S1** summarizes the six different strategies used in this study.

It is worth noting that although NiMARE could natively support any kind of data masker for both volumetric and surface data, it relies on Nilearn’s masker (Abraham et al., 2014), which does not currently support the transformation from surface to volume space. Therefore, we implemented a surface version of `CorrelationDecoder` to perform the functional decoding on surface space, given that gradient methods are natively implemented on the cortical surface (Vos de Wael et al., 2020). First, we transformed meta-analytic maps across decoding strategies from the MNI152 space to the standard MNI fsLR 32K 2-mm mesh surface space of the HCP, using the `mni152_to_fslr` function from the neuromaps’ transforms module (Markello et al., 2022). Then, similar to the `CorrelationDecoder` approach, we generated correlation coefficients between vertex-level unthresholded meta-analytic maps and unthresholded pseudo-activation maps from the gradient segments. As described in Step 2.2 (**Fig. 1, Step 2.2**), *n* = 31 different segmentations of the principal gradient were generated, corresponding to numbers of segments ranging from *k* = 2 to *k* = 32.

#### Evaluating Significance of Correlation Values

Note that `CorrelationDecoder` does not test if null hypotheses of the correlations are driven by the specific topography of the map of interest. To address this issue, permutation tests have been implemented (Alexander-Bloch et al., 2013a, 2013b; Vandekar et al., 2015), comparing the empirical correlation between two spatial maps of interest to a set of null correlations. We used a permutation test to evaluate the null hypothesis that the meta-analytic and gradient maps were not significantly different. The null correlations were generated by correlating the maps of interest with a group of null maps. We leveraged non-parametric spatial permutation models, also known as the “spin permutation” model (Alexander-Bloch et al., 2018; Baum et al., 2020; Cornblath et al., 2020; Váša et al., 2018; Vázquez-Rodríguez et al., 2019). We generated 1000 null maps with identical spatial autocorrelation to the maps of interest in surface space by randomly rotating the spherical projection of the meta-analytic maps. In particular, we applied the Váša method (Váša et al., 2018) using the `gen_spinsamples` function from the Python netneurotools’s stats module (Markello and Misic, 2021). This method produces a lower false positive rate than naive, parameterized, and non-parameterized data models; it also generates permutations without duplicates in the parcel assignments and without rotating the medial wall (Markello and Misic, 2021). Non-parametric p-values were estimated from the null distribution of correlation values by calculating the fraction of null maps that generated a correlation equal to or greater than the correlation of interest. Finally, a false discovery rate (FDR) correction was performed using the Benjamini-Hochberg procedure (Benjamini and Hochberg, 1995). We considered the correlation between meta-analytic maps and gradient maps significant for FDR-corrected p-value < 0.05.

### Analysis Step 4: Performance of Decoding Strategies

Next, we evaluated the six decoding strategies for each segmentation approach (i.e., 18 pipelines). We used multiple metrics to compare the relative performance and utility across decoding strategies (**Table 1**). Our goal was to identify a decoding strategy with a high association with the gradient maps and meaningful and informative terms associated with coherent cognitive functions. Subsequent sections describe each metric in detail.

**Table 1:**
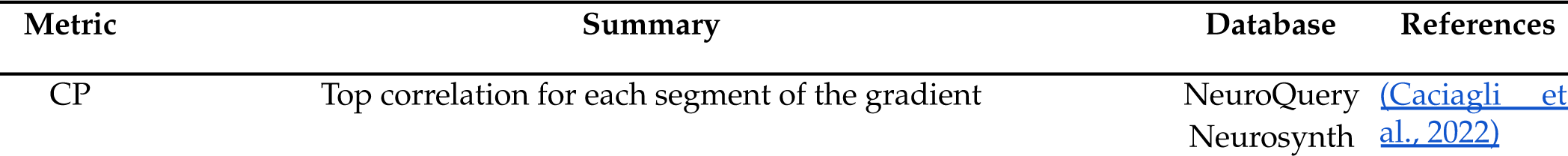
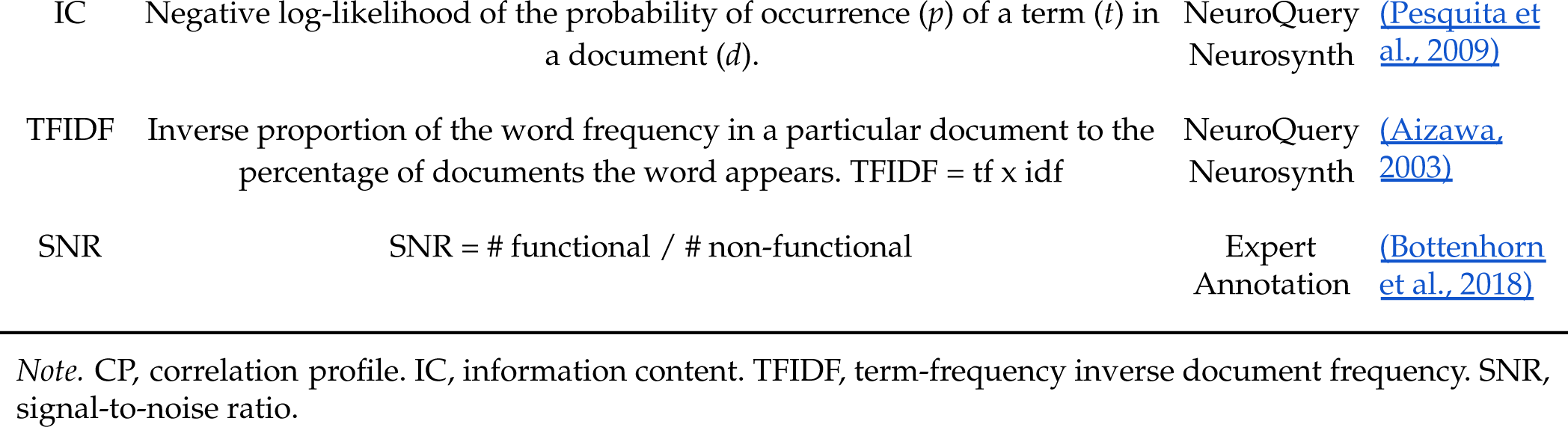
Summary of Decoding Performance Metrics.

#### Meta-Analytic Correlation Profile

A successful gradient decoding framework should be able to produce meta-analytic maps with a high correlation across segments of the gradient (**Fig. 1, Step 4.1**). It is common practice in continuous decoding approaches to select the term/topic maps that show the highest correlation with the target map and use the associated labels to describe the input map. We selected the meta-analytic map with the maximum correlation value for each segment for further analysis. We defined a correlation profile for the principal gradient as a two-dimensional representation of the gradient axis, where the x-axis corresponds to the coordinates in gradient space and the y-axis corresponds to the top correlation for each gradient segment. The correlation profile metric was inspired by previous work (Caciagli et al., 2022). Given the large number of evaluated meta-analytics maps, these correlations could be high by design. In addition, the different number of meta-analytic maps per strategy prevents us from comparing the correlation values that differ from the maximum.

#### Semantic Similarity

A successful decoding framework should also be able to classify a map of interest with meaningful and informative terms. To this end, we used two different semantic similarity node-based metrics (**Fig. 1, Step 4.2**) to evaluate the properties of terms (Pesquita et al., 2009).

First, we examined the information content (IC), which measures how specific and informative a term is based on its annotation frequency. The IC of term *t* in document *d* is defined as the negative log-likelihood of the probability of occurrence (*p*) of a term in a document:

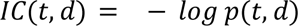

Higher values of IC are associated with less probable terms and thus more specific concepts (e.g., pain). Lower values correspond with high probable terms and thus more general/abstract concepts (e.g., fMRI, resonance) (Zhu and Iglesias, 2017). We used the combined corpora of article abstracts from Neurosynth and article abstracts, bodies, author-provided keywords, and titles from NeuroQuery to calculate the probability of occurrence *p*(*t, d*) = *f*(*t, d*)/*N*(*d*), where *f*(*t, d*) is the number of times the term *t* appears in document *d* and *N*(*d*) is the total number of terms from the vocabulary in document *d*, *N*(*d*) = Σ *f*(*t*’, *d*). Next, the IC value of a term associated with a meta-analytic map was etermined by the average across the documents included in the meta-analysis. In the case of topic-based meta-analytic maps, the IC was calculated as the sum of individual ICs from the top words in the topic.

Second, we computed an additional metric that accounts for the popularity of a term, thus establishing a balance between the specificity (i.e., IC) and the popularity (e.g., frequency of annotation) of a term (Aizawa, 2003). This metric is known as the term frequency-inverse document frequency (TFIDF), which is defined as the proportion of the frequency of the word in a particular document (term frequency: *tf*) to the percentage of documents the word appears in (inverse document frequency: *idf*):

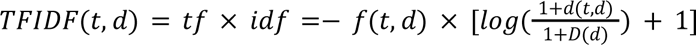

where *tf* is the number of times that the term *t* appears in the abstract of document *d f*(*t, d*), and *idf*(*t, d*) is the negative log-likelihood of the proportion of the number of documents where the term appears to the total number of documents in the corpus *D*. The constant “1” was added to the denominator and denominator to prevent zero division, and another “1” was added to the logarithm to avoid ignoring terms with zero idf. The `TfidfTransformer` class from the Python library `scikit-learn` (Abraham et al., 2014) was used to calculate the TFIDF values. TFIDF determines how relevant a given word is in a particular document. Common words in a single or a small group of documents tend to have higher TFIDF numbers than common words, such as articles and prepositions. The term’s relevance in a meta-analytic map can be computed, similarly to the IC, by averaging across the studies used in the meta-analysis. Moreover, the TFIDF of a topic-based decoder was calculated as the sum of individual TFIDF from the top words in the topic.

#### Signal-to-Noise Ratio Profile

Finally, in a successful gradient decoding framework, the top terms of a particular segment of the gradient axis should be associated with clear, coherent, and useful cognitive terms, facilitating meaningful functional interpretations. However, Neurosynth terms are automatically extracted from papers, and not all extracted terms contain helpful information. That is, terms such as “*memory*” or “*attention*” carry important functional meanings and are thus more useful than “*brain*” or “*association*”. Therefore, in a typical decoding workflow, researchers often manually filter meta-analytic terms from Neurosynth and decode maps using the most meaningful terms, a process that a single individual commonly carries out. We used Neurosynth and NeuroQuery terms to calculate each decoding framework’s signal-to-noise ratio (SNR) (**Fig. 1, Step 4.3**).

For Neurosynth terms, we implemented a crowdsourced approach in which ten neuroimaging researchers of varying backgrounds and career stages provided manual annotations of meta-analytic terms. Each term was classified according to four categories: “*Anatomical*”, “*Functional*”, “*Clinical*”, and “*Non-specific*”; these categories were initially described by Bottenhorn et al. (Bottenhorn et al., 2018). “*Anatomical*” terms describe regions, networks, or locations in the brain (e.g., “cortex”, “acc”, “mpfc”). “*Functional*” terms describe mental processes, tasks, or behaviors (e.g., “hand movements”, “visual”). “*Clinical*” terms describe participant-related characteristics, including clinical diagnoses and symptoms (e.g., “adhd”, “alzheimer”), as well as age and gender. “*Non-specific*” terms provide no useful information about the study, such as general terms found in articles (e.g., “useful”, “influences”) or methodological terms (e.g., “correlation”, “high resolution”). For NeuroQuery, we also used the expert annotation for terms present in Neurosynth. The rest of the terms were annotated based on the classification provided by NeuroQuery, which is available on GitHub (www.github.com/neuroquery/ neuroquery_data/blob/main/data/data-neuroquery_version-1_termcategories.csv).

Using the annotations, we created a frequentist probability distribution to classify each term automatically: *p(category|word)*. We classified a term using the category with a probability greater than 0.5. For topic-based decoding, we used the term classification distribution to classify topics. In this case, the probability for topics was calculated as the sum per category of the *p(category|word)* weighted by the *p(word|topic)*. If the categories failed to reach the 0.5 thresholds, the topic was classified as “*Non-specific*”. As demonstrated by Bottenhorn et al., SNR for a decoding framework is defined as the proportion of functional (i.e., “*Functional*”) to non-functional (i.e., “*Anatomical*”, “*Clinical*”, and “*Non-specific*”) terms (Bottenhorn et al., 2018). The SNR for each segment solution of the connectivity gradients was calculated from the classification of the individual meta-analytic maps associated with each spectrum segment. Greater SNR was expected for a more informative functional characterization of the connectivity gradient.

#### Visualization of Decoded Gradient Maps

The final step of meta-analytic decoding is to report findings and generate an appealing and effective visualization. Across the literature, two different visualization strategies are commonly employed. First, radar plots (Bartley et al., 2019; Chang et al., 2013; Chase et al., 2020; de la Vega et al., 2016) illustrate the top meta-analytic terms or topics associated with a decoded map. Second, word clouds (Fan et al., 2022; Laubach et al., 2018; Riedel et al., 2019; Rubin et al., 2017) represent term frequency. We combined the two plots in a single figure to show a hierarchical view of the decoding results across gradient maps (**Fig. 1, Step 5**). Before generating the visualizations, we filtered out the meta-analytic terms/topics without coherent cognitive function association using our expert annotations (i.e., “*Non-specific*” terms were excluded) and those with correlations that were not statistically significant.

### Analysis Step 5: Multidimensional Decoding

Finally, we propose methods for decoding lower-order gradient maps (e.g., second and third gradients). Naturally, we could consider applying the same segmentation and decoding approach proposed in the last four analysis steps to lower-order gradients separately. The second and third gradients normally account for around 15% and 10% of the variance, respectively; thus, they still represent significant connectivity information. However, decoding these maps alone could mislead functional interpretations, given that the new segments for decoding would be defined with limited information from the original connectivity matrix.

Therefore, we suggest analyzing all the components together in a high-dimensional space. As was shown in the work by Margulies and colleagues, analysis of the first and second gradient gradient together yielded results in which the additional second component separated one end of the spectrum in the somatomotor and visual networks (Margulies et al., 2016). Similarly, adding the third component to the analysis can delineate the frontoparietal network in the intermediate transition from transmodal to unimodal regions within the gradient (Smallwood et al., 2021). In this high-dimensional space, the joint gradient will account for more than 70% of the variance, improving the performance of the segmentation algorithm given that a high-dimensional space will provide a better approximation of the n-dimensional distances from the original connectivity matrix.

In this high-dimensional space, the clusters are defined only by the k-means approach. This method is the only approach from the previous analysis compatible with high-dimensional data. The best solution for the number of components and number of clusters was selected based on the same criteria from the previous analysis using the mean silhouette coefficient (Rousseeuw, 1987), the variance ratio score (Caliński and Harabasz, 1974), and the cluster separation score (Davies and Bouldin, 1979). Then, the pseudo-activation maps were defined with the same Gaussian kernel described above and calculated with respect to the center of inertia within each cluster.

## Results

### Analysis Step 1: Functional Connectivity Gradient

To achieve the goals of this paper, we used the rs-fMRI group-average dense connectome from the Human Connectome Project (HCP) S1200 data release. The z-values from the dense connectome were inverted to correlation coefficients, thresholded, and converted to cosine similarity values. This matrix comprises 91,282 X 91,282 grayordinates, including cortical regions (59,412 vertices) in surface space and subcortical structures (31,870 voxels) in volume space. Diffusion embedding was applied to the cosine similarity affinity matrix using the `mapalign` repository. We used the gradient with the highest variance explained, the principal gradient, for the first four analyses. Only the cortical regions were considered for this analysis. We added the 5,572 vertices from the medial wall for 64,984 vertices (32,492 per hemisphere) for visualization. **Fig. 1S** shows the connectivity matrix, affinity matrices, and the principal gradient.

### Analysis Step 2: Segmentation and Gradient Maps

Our first aim was to evaluate three segmentation approaches: (i) percentile-based segmentation, (ii) segmentation based on a 1D *k*-means clustering approach, and (iii) segmentation based on the Kernel Density Estimation (KDE) curve of the gradient axis. For each segmentation approach, *n* = 31 different segmentations of the principal gradient were generated, corresponding to segments ranging from *k* = 2 to *k* = 32, for a total of 93 maps. **Fig. 2** shows the segmentation for 3, 17, and 32 segment solutions. **Fig. 2a** shows the position of the boundaries as determined by the different segmentation algorithms, with the peak activation point (red dot), which corresponded to the median point, center of inertia, and local maximum for PCT, KMeans, and KDE, respectively. We notice more diverse boundaries across segmentation methods for small-segment solutions, with KMeans and KDE showing overlapping solutions at the right end of the spectrum. In contrast, the boundaries determined by PCT and KMenas for a large-segment solution show some overlapping. In **Fig. S5,** we reported the similarity between the three segmentation methods across segmentation solutions. The separations provided by the segmentation approaches were more similar for a large number of segment solutions. Next, we determined pseudo-activation maps using a Gaussian kernel of the Euclidean distances defined in the gradient space, minimizing the effect of boundaries. These maps were the target of the decoder algorithms. **Fig. 2b** illustrates the resultant continuous pseudo-activation maps by segments.

**Figure 2.**
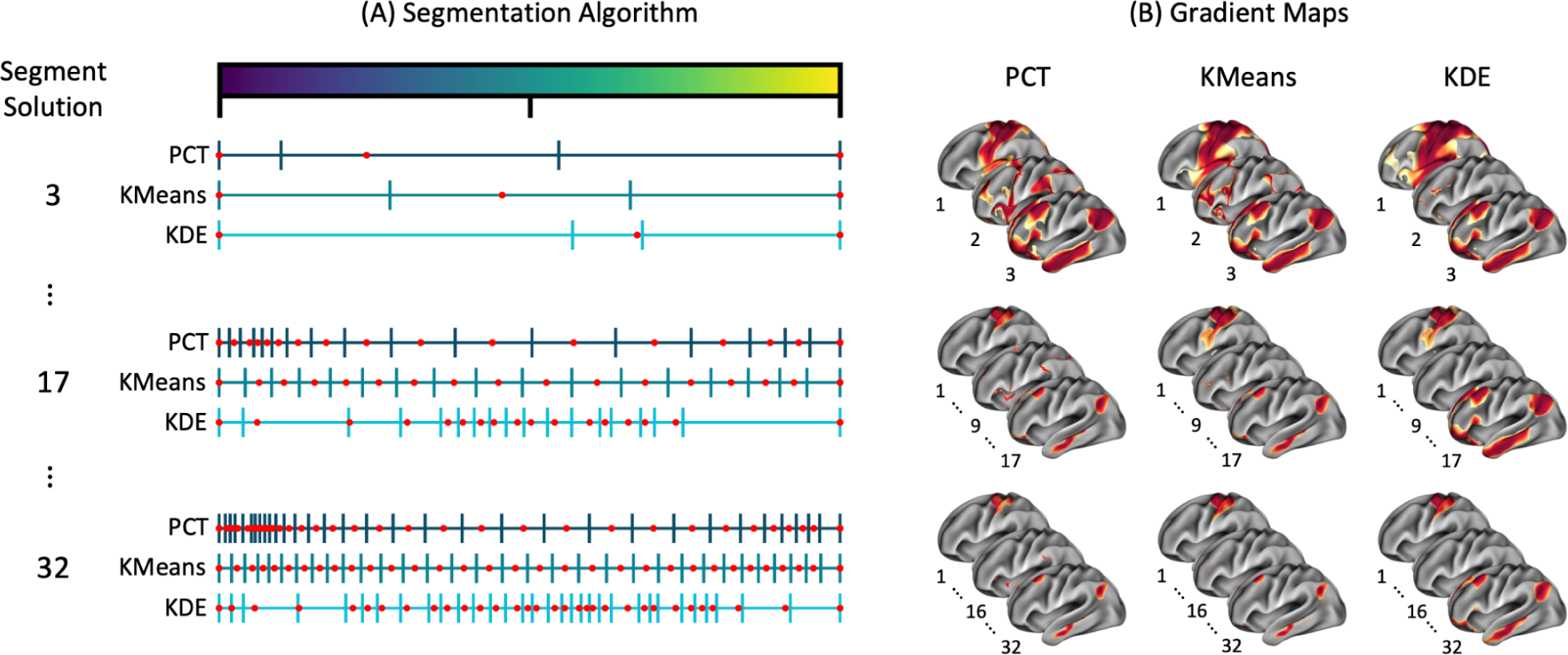
Segmentation of the Principal Gradient. Segmentation of the principal gradient with three different approaches: PCT, KMeans, and KDE. 31 different segmentations were generated, ranging from a solution of 2 to 32 segments. A) Three distinctive and representative solutions are provided for 3, 17, and 32 segments. The segmentation algorithm determines the boundaries, where the red point indicates the peak activation corresponding to the median point, center of inertia, and local maximum for PCT, KMeans, and KDE, respectively. B) The continuous vertex-level values range between 0 and 1, where 1 represents values closer in space or strongly connected to the reference point, and 0 indicates elements closer to the boundaries or weakly connected to the reference point. See **Fig. S4** for the complete set of segment solution plots. The rest of the gradient map plots are in the Supplementary Results.

Silhouette scores were computed for each segment solution and segmentation algorithm at the vertex level. **Fig. 3** presents a comprehensive analysis of the silhouette coefficient by depicting it in three distinct ways: a) as a distribution, b) in terms of cluster imbalance, and c) through confidence maps. **Fig. 3a** illustrates the distribution of the silhouette coefficient for three different clustering algorithms (i.e., PCT, KMeans, and KDE) and for three distinctive segment solutions. On average, the three methods performed best for a small segment solution, with KMeans slightly better than the others, with a median value over 0.2. PCT and KDE showed the highest number of vertices with negative silhouette coefficients, indicating vertices assigned to the wrong segment. The maximum silhouette score observed for this solution was around 0.4. However, with an increase in the number of segments, the average performance of all methods declined for elements inside intermediate segments and improved for elements inside the two end segments. For example, for the 17-segment solution, KMeans and KDE showed a small group of elements with values over 0.6 silhouette score. Interestingly, the KDE method shows a higher concentration of vertices with positive silhouette coefficients, indicating a better clustering solution, given that most of the local maximum were in the middle of the spectrum.

**Figure 3.**
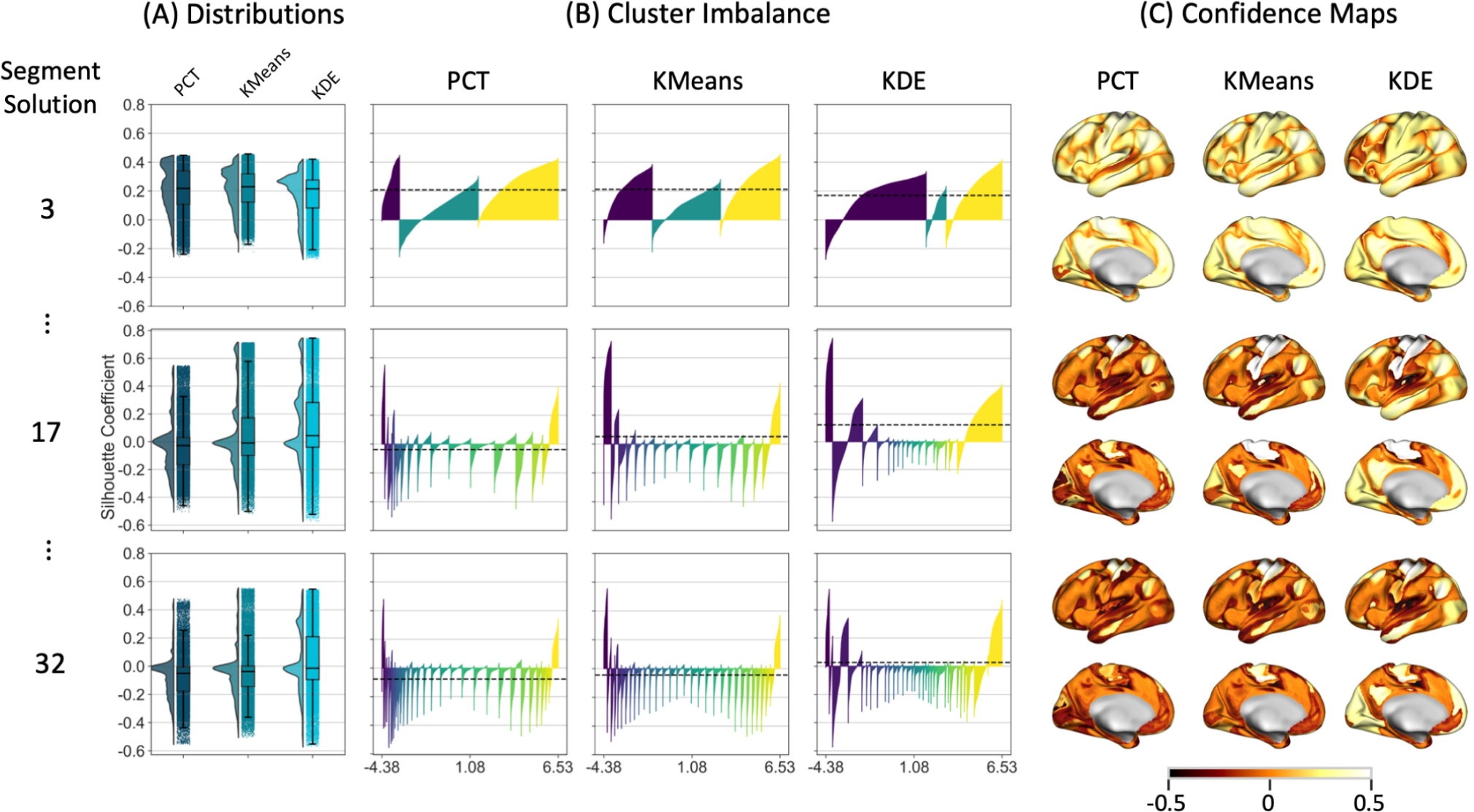
Silhouette Coefficients, Cluster Imbalance, and Confidence Maps. A) Distributions of vertex-wise silhouette coefficient are provided for three different and representative segment solutions (e.g., 3, 17, and 32 segments); the values range from -1 to 1, where +1 indicates that the vertex was assigned to the correct cluster, a value of 0 indicates that the vertex is on or very close to the decision boundary between two neighboring clusters, and -1 indicates the vertex might have been assigned to the wrong cluster. B) Cluster imbalance is plotted as the silhouette coefficient density for each segment solution and segmentation approach per segment to visualize the density of vertices that were assigned to the correct (i.e., positive values) or the wrong (i.e., negative values) segments within each segment. Within each segment, values in the x-axis are sorted according to their values in the y-axis. C) Confidence maps are represented by the silhouette coefficient of each vertex to its assigned network. Regions close to the boundaries between networks showed less confidence in their assignment. The best value is 1, and the worst is -1; values near 0 indicate overlapping clusters. The distribution, cluster imbalance, and confidence maps for the rest of the segment solutions can be found in the Supplementary Results.

**Fig. 3b** presents the silhouette coefficient across segments within each segment solution. For a small number of segments (e.g., three segments), the PCT solution yielded segments at both ends of the spectrum with all positive coefficients; nevertheless, it wrongly assigned a significant number of vertices to the intermediate segment. KDE yielded a small fraction of segments incorrectly assigned in the first and last segments. Conversely, KMeans shows the most precise assignment with a more balanced number of vertices with positive silhouette coefficients across segments. Increasing the segment solution diminished the performance of the three methods, especially in intermediate segments. However, the silhouette coefficient increased in segments at one end of the gradient axis. For example, for the 17-segment solution, KMeans and KDE showed values over 0.6 silhouette score at the left-side end. In general, the highest balance in assignment across segment solutions was produced by KMeans. Lastly, **Fig. 3c** illustrates which regions of the cortical surface were most affected by the segmentation approach. Align with Fig 3b, the values with the highest confidence were found in the segments located at both ends of the spectrum.

Mean silhouette scores, variance ratio, and cluster separation were computed to determine relative performance across percentile-based, k-means-based, and KDE-based segmentations. The highest mean silhouette score, highest variance ratio, and lowest cluster separation represented the best performance. As illustrated in **Fig**. **4**, KDE exhibited the highest average of the mean silhouette coefficient across different segment solutions, ranging between 0.05 and 0.15 for 5-32 segment solutions. The variance ratio of the three methods was very similar across segment solutions. In contrast, PCT showed the lowest cluster separation across segment solutions. Overall, all segmentation approaches yielded better results for two segment solutions. The best scores were silhouette = 0.24 (KMeans), variance ratio = 28575 (PCT), and cluster separation = 1.29 (KDE). In particular, KMeans achieved the best top two performances across the three metrics (silhouette = 0.24, variance ratio = 28566, cluster separation = 1.32) for the two-segment solution (**Fig. S6**). These findings highlight the importance of choosing the most appropriate clustering method based on the number of segments required and the desired accuracy.

**Figure 4.**
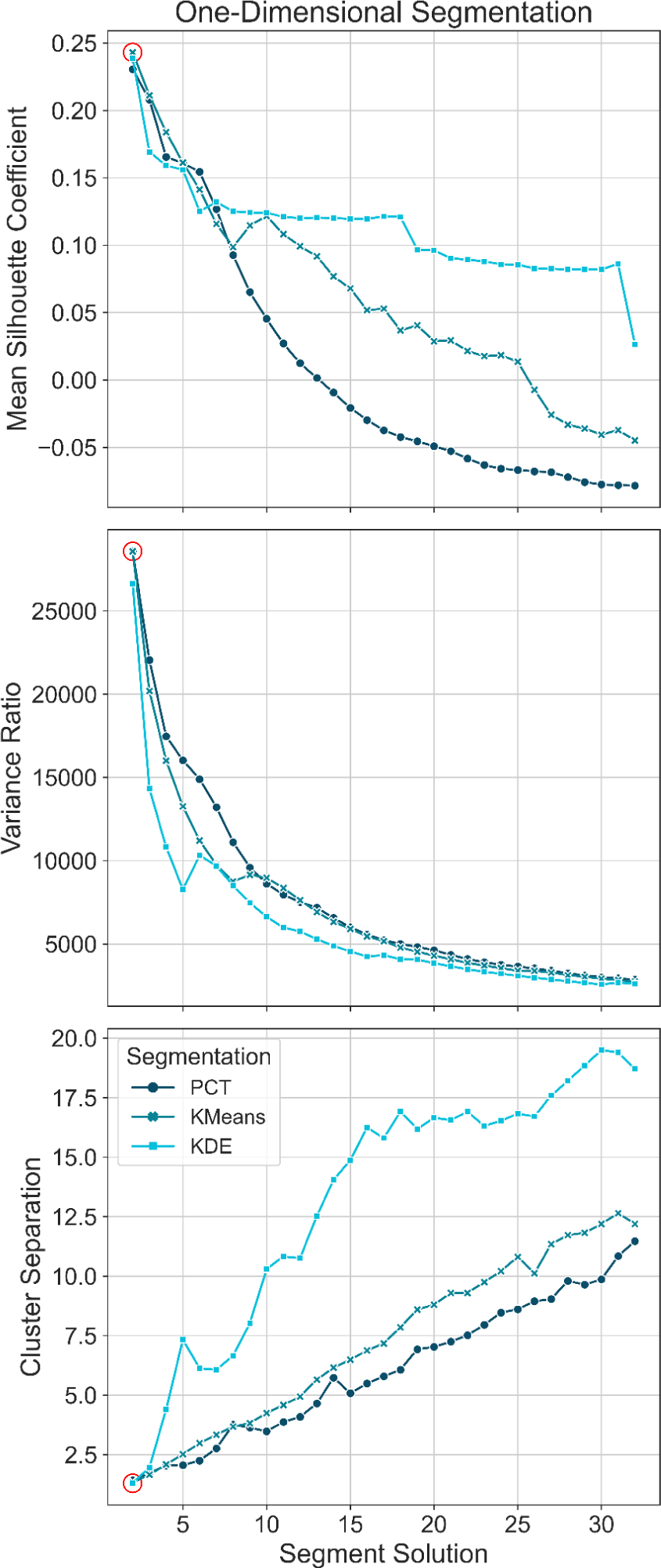
Mean Silhouette, Variance Ratio, and Cluster Separation Scores. Mean silhouette coefficient, variance ratio, and cluster separation plotted by segment solution across segmentation approaches to determine relative performance, with the highest mean silhouette score, highest variance ratio, and lowest cluster separation representing the best performance (red circle).

### Analysis Step 3: Meta-Analytic Functional Decoding

Next, six different sets of meta-analytics maps were produced by the combination of two meta-analytic databases (i.e., Neurosynth and NeuroQuery) and three meta-analytic methods (i.e., term-based, LDA-based and GC-LDA-based meta-analysis). The mata-analytics maps that resulted from each strategy can be found in the Supplementary Results (**Fig. S7**).

### Analysis Step 4: Performance of Decoding Strategies

Our second aim was to assess the performance of 18 meta-analytic decoding/segmentation strategies using multiple metrics, including a correlation profile, two semantic similarity measures, and a normalized signal-to-noise ratio (SNR).

#### Correlation Profile

First, we assumed that a successful gradient decoding framework should be able to produce meta-analytic maps with a high correlation across segments of the gradient. Thus, we next performed a detailed analysis of the correlation profile and maximum correlation coefficient, as shown in **Fig. 5**. The correlation profile curve (**Fig. 5a**) shows an exponential decrease in the average of maximum correlation coefficients across segment solutions for each combination of segmentation approach, meta-analytic method, and database. Then, the correlation curve reaches a plateau for segment solutions with more than 15 segments. The highest average correlation value was the two-segment solution of the NQ-TERM-PCT approach (0.73 ± 0.03). The best performance across segment solutions was achieved through PCT segmentation and term-based decoder with Neurosynth (0.23 ± 0.14). GC-LDA-based decoders showed the worst performance across segmentation solutions (0.16 ± 0.15). In general, we did not observe a considerable difference between NS and NQ combinations. In some cases, NS performed slightly better (e.g., 17-segment solution: NS = 0.19 ± 0.16, NQ = 0.18 ± 0.16), and in others, NQ showed higher correlation values (e.g., three-segment solution: NS = 0.46 ± 0.22, NQ = 0.47 ± 0.22). Concerning segmentation approaches, PCT performed best (0.21 ± 0.13), while KMeans (0.18 ± 0.16) performed better than KDE (0.15 ± 0.19) across segmentation solutions. **Fig 8a** provides a complete comparison, including all models and segment solutions.

**Figure 5.**
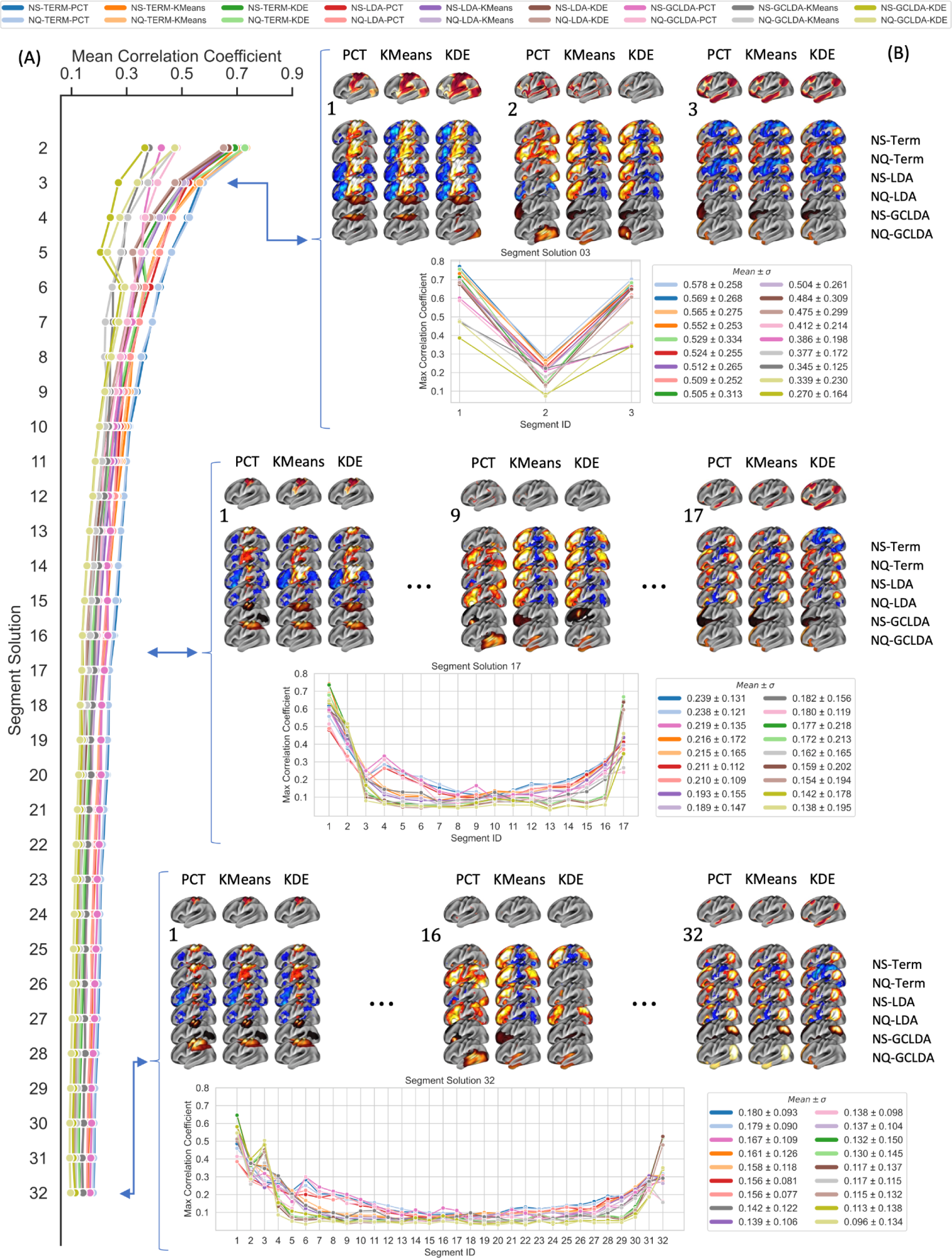
Correlation Profile. Correlation profile of the means within segment solution of maximum correlation coefficients across the combination of models (Term, LDA, and GC-LDA), databases (Neurosynth: NS and Neuroquery: NQ), and segmentation approach (PCT, KMeans, and KDE). In the horizontal block, three distinctive segment solutions (3, 17, and 32) are shown, containing the meta-analytic maps (the second 6 rows) that showed the highest correlation coefficient with gradient maps (top row) from a segmentation approach (columns). The horizontal plot shows the correlation profile within a segment solution, and the rectangular box at the right shows each model’s mean and standard deviation of each model within each segment solution. The correlation profile for the rest of the segment solutions can be found in the Supplementary Results.

**Fig. 5b** represents the correlation profile within segment solutions. The profile exhibits a u-shaped curve, where correlation at both ends of the spectrum was higher than the rest of the segments, with the left end showing the highest correlation coefficient from maximum values around 0.7 for three segments to around 0.6 for 32 segments. Similarly, the right end shows maximum values of 0.6 and 0.3 for three and 32 segments, respectively. The low correlations achieved by all models in the middle of the spectrum drove the low overall performance of all decoder approaches. An increase in the segment solution also increased the number of intermediate segments, making it worse. The decrease of the standard deviation of the mean with the increase in the segment solution also showed a rise in most segments with low performance in the overall mean per segment solution. Overall, including additional low-correlation values from intermediate segments drove the decrease in overall performance with the increase in segment solution. In particular, we observed a change in the correlation coefficient with the size of the segments for maps with less than 10,000 vertices (**Fig. S8**). These vertices represent approximately 15% of the brain cortical surface on MNI fsLR 32K.

#### Semantic Similarity

Second, we assumed that a successful decoding framework should produce meaningful and informative terms. Therefore, we used two different semantic similarity node-based metrics to evaluate properties of words. For example, information content (IC), which measures how specific and informative words are based on their annotation frequency, and frequency-inverse document frequency (TFIDF), which accounts for the term’s popularity in the whole corpus. **Fig. 6** summarizes the results of these semantic similarity metrics across segment solutions and for three different segment solutions.

**Figure 6.**
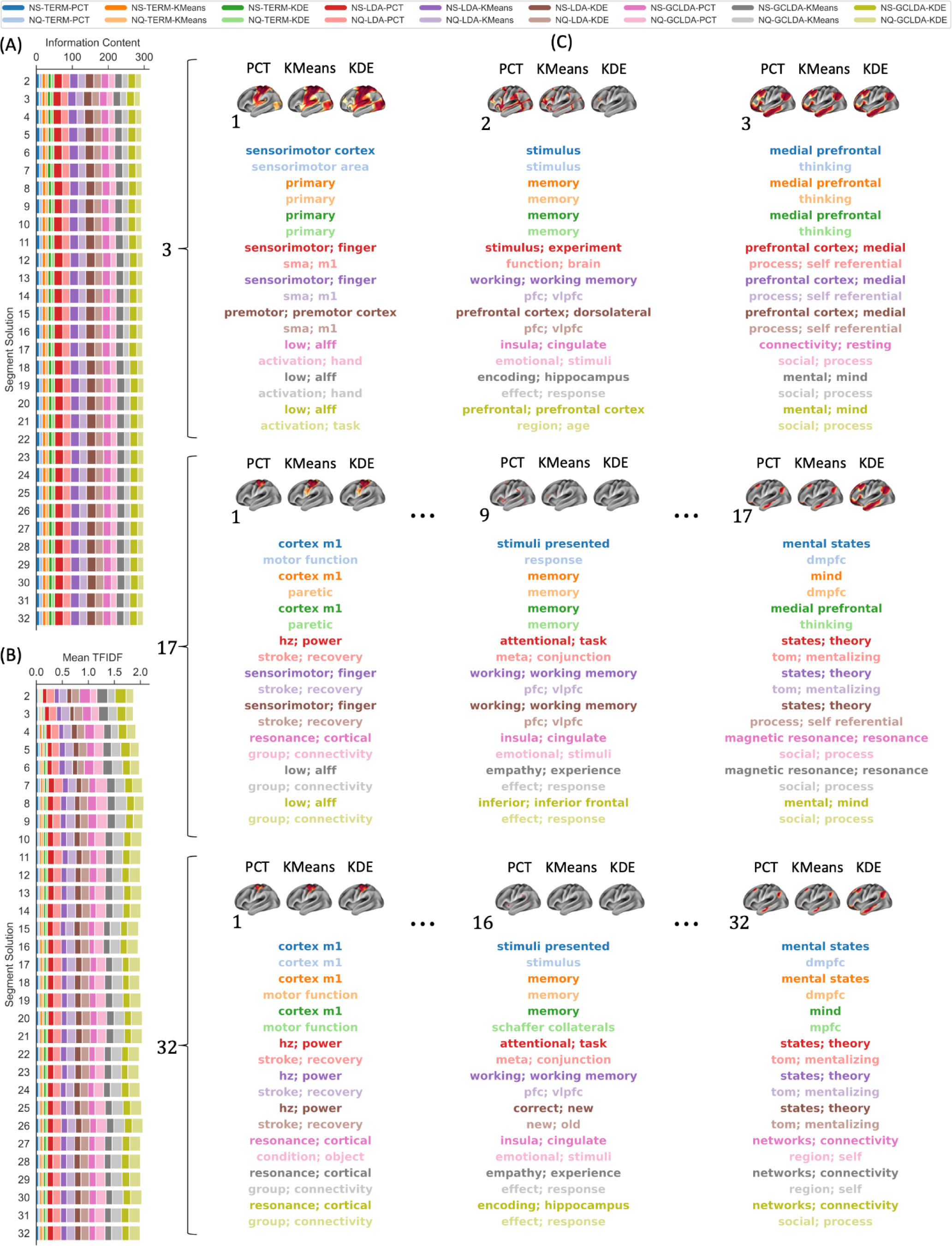
Semantic Similarity Profile. Average semantic similarity metrics: A) information content (IC) and B) term-frequency inverse document frequency (TFIDF) across segment solutions for the 18 different decoding strategies. C) Terms associated with the meta-analytic maps of maximum correlation with the corresponding segment ID. The IC and TFIDF profile for each segment solution can be found in the Supplementary Results.

**Fig. 6a** presents the result of the mean information content of the top term across segment solutions. Decoding strategies that included LDA or GC-LDA yielded a collection of terms that resulted in the highest information content, with LDA (IC = 22.40 ± 3.21) showing a slightly better performance than GC-LDA (IC = 18.57 ± 3.18). Moreover, the topic-based strategies that used NS (IC = 17.87 ± 7.22) performed better than those trained with NQ (IC = 15.11 ± 5.49). On the other hand, term-based models only produced a single word per map, which yielded the worst information content (IC = 8.48 ± 2.30) of all pipelines. **Fig. 6b** illustrates the mean TFIDF of the top terms for all segment solutions. Similarly, topic-based models produced relatively good performance (LDA-TFIDF = 0.13 ± 0.06, GC-LDA-TFIDF = 0.17 ± 0.08), with the model that uses NQ (TFIDF = 0.13 ± 0.09) having higher TFIDF than NS (0.09 ± 0.07) across segment solutions. Term-based decoders also performed poorly (TFIDF = 0.04 ± 0.02) for this metric. **Fig 8a** provides a complete comparison, including all models and segment solutions.

In **Fig. 6c**, we present the top words produced by the different decoding strategies for three distinctive segment solutions and three subsegments within each segmentation solution. For solutions with a small number of segments, we observed a high discrepancy between the top words between the models. However, within each model, we noticed a certain similarity in the terms, for example, “sensorimotor cortex” and “primary” for the left end of term-based decoders, “sensorimotor” and “sma” for LDA, and “activation” and “low” for GC-LDA. In particular, strategies where only the segmentation approach was different show the highest similarities. Similarly, for larger-segment solutions (e.g., 17 and 32), the words within the meta-analytic decoder continued to be very similar, and there was still a more substantial difference between modes. Interestingly, the terms at both ends for this number of segments were associated with terms with less cognitive content, for example, words like “cortex”, “cortical”, “magnetic”, and “resonance”, which do not provide much value for functional interpretation of a given map.

#### Signal-to-Noise Ratio

Finally, we assumed that in a successful gradient decoding framework, the top terms that label a particular segment of the gradient axis should be associated with clear, coherent, and valuable cognitive terms to facilitate meaningful functional interpretations. For that purpose, terms were classified according to four categories: “*Anatomical*”, “*Functional*”, “*Clinical*”, and “*Non-specific*” by a team of ten neuroimaging researchers. In particular, we were interested in functional terms. **Fig. S9** presents the result of the classification.

We next calculated the normalized signal-to-noise ratio (SNR) for each decoding strategy using the classification of the top terms across subsegments for each segment solution. Here, the SNR for a decoding framework was defined as the normalized proportion of functional to non-functional terms. **Fig. 7** presents the results of the SNR across segment solutions and the individual classification within three distinctive solutions. **Fig. 7a** illustrates the normalized SNR values of all decoding frameworks. Term- and GCLDA-based decoders showed the highest SNR values for all segment solutions, while LDA decoders produced the lowest SNR across segment solutions. Regarding meta-analytic databases, NQ showed the highest SNR for a two-segment solution. See the heatmap in **Fig 8a** for a complete comparison, including all models and segment solutions.

**Figure 7.**
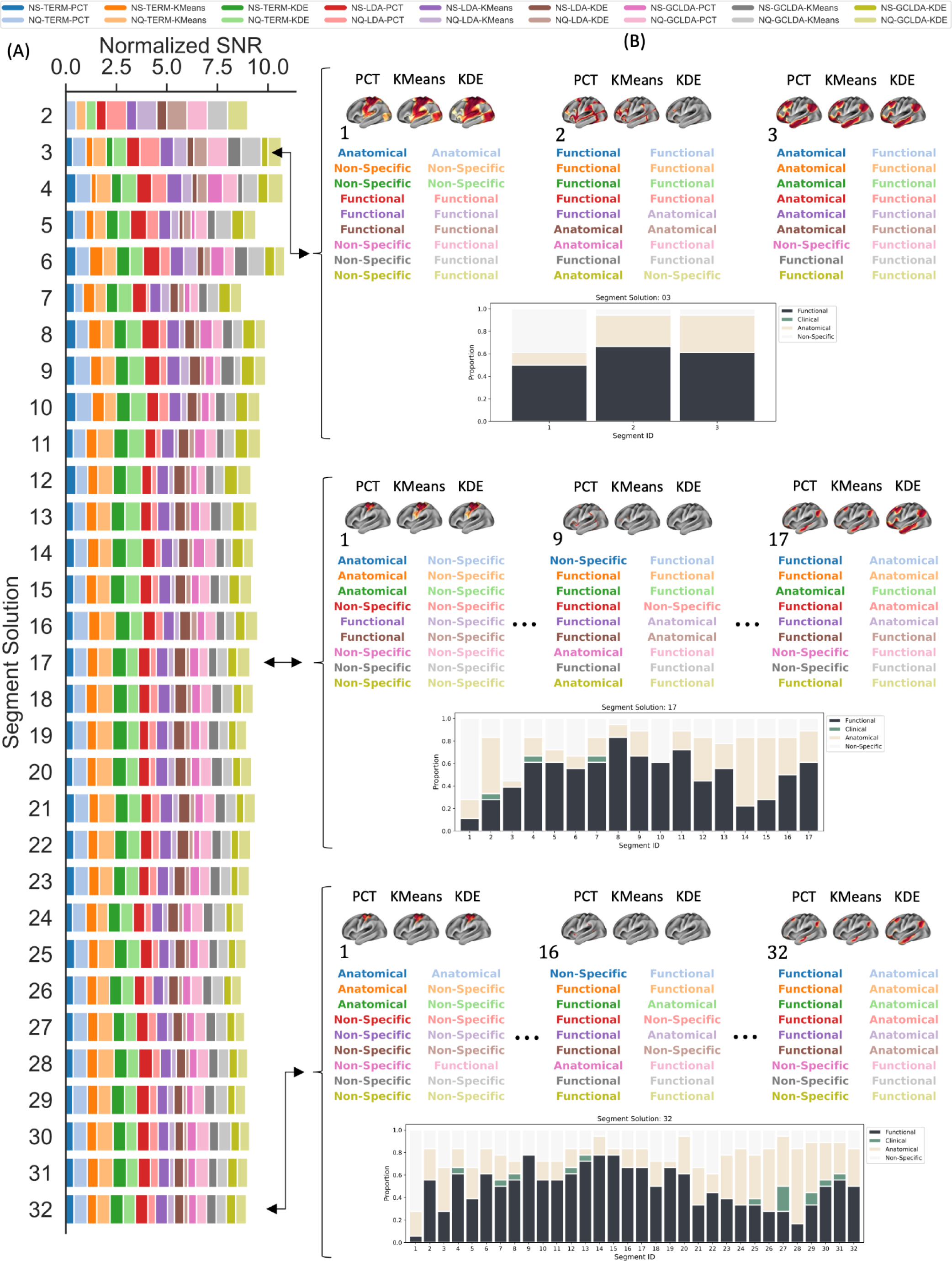
Normalized Signal-to-Noise Ratio. A) SNR from classifying terms and topics across segment solutions for the 18 decoding strategies. B) Classification of the term associated with the meta-analytic maps of maximum correlation with the corresponding segment ID; and proportion of “*Functional*”, “*Clinical*”, “*Anatomical*”, and “*Non-specific*” terms within segment solution. The classification plots for the rest of the segment solutions can be found in the Supplementary Results.

**Fig. 7b** presents the individual classification within three distinctive segment solutions. For a small number of segments, we see that most models found a functional term/topic at the end of the spectrum (i.e., 50 and 60 for the left and rightmost segments, respectively). For intermediate segments, slightly over 60 percent of the terms were classified as functional. However, an increase in the number of segments yielded fewer functional terms at both ends of the spectrum while producing more functional terms in middle segments. Generally, frameworks that used NQ were most likely associated with functional terms at both ends of the spectrum, regardless of the meta-analytic model or segmentation approach. For a large number of segments (e.g., 32), NS did not produce any top maps at the left end that were associated with functional terms, as they were classified as “*Anatomical*” or “*Non-Specific*”. In particular, we noticed that when the term was classified as non-functional, terms tended to be associated with “*Anatomical*” or “*Clinical*” words. In contrast, LDA- and GC-LDA-based decoders were mainly associated with “*Non-specific*”. Lastly, we noticed that the difference in the functional terms between NS and NQ tends to be less distinctive in the middle of the spectrum.

We next sought to filter the results of the top meta-analytic maps and only keep those classified as “*Functional*”. After examining the results from all the functional decoders together, it was clear that the segmentation technique had minimal to no influence on the ultimate result of functional decoding for a large number of segment solutions at both ends of the spectrum, as the word clouds showed similar words regardless of the segmentation algorithm used (**Fig. S10**). The stability in clustering at both ends of the gradient, especially for a large-segment solution (**Fig. S5**), results from applying a data-driven segmentation approach (e.g., PCT, Kmeans, or KDE) to a 1D axis (i.e., the principal gradient).

#### Overall Performance and Optimal Strategy

Together with the above results, we reviewed 18 combinations of segmentation approaches, meta-analytic decoders, and databases. **Fig. 8** presents the results of the four benchmark metrics for the 18 methods and 31 segment solutions, along with a heatmap summarizing the overall performance. In **Fig. 8a**, we marked the strategies that performed over the threshold with a red square. We defined the threshold using a 90^th^ percentile for the correlation coefficient and SNR and a 70^th^ percentile for IC and TFIDF, which were the maximum values, respectively, that yielded a unique solution for the overall performance. In **Fig. 8b**, results shown in the five heatmaps from **Fig 8a** are summarized. The best-performing strategy was the LDA-based meta-analysis on the NeuroQuery database applied to a two-segment solution determined by the k-means algorithms (NQ-LDA-KMeans for two-segment solution). The only metric where NQ-LDA-KMeans did not perform well was SNR, which can be addressed in the future by filtering non-functional terms from the decoding model. Here, we also accounted for the performance of the segmentation approaches alone, as determined in analysis step 2 (**Fig. S6**).

**Figure 8.**
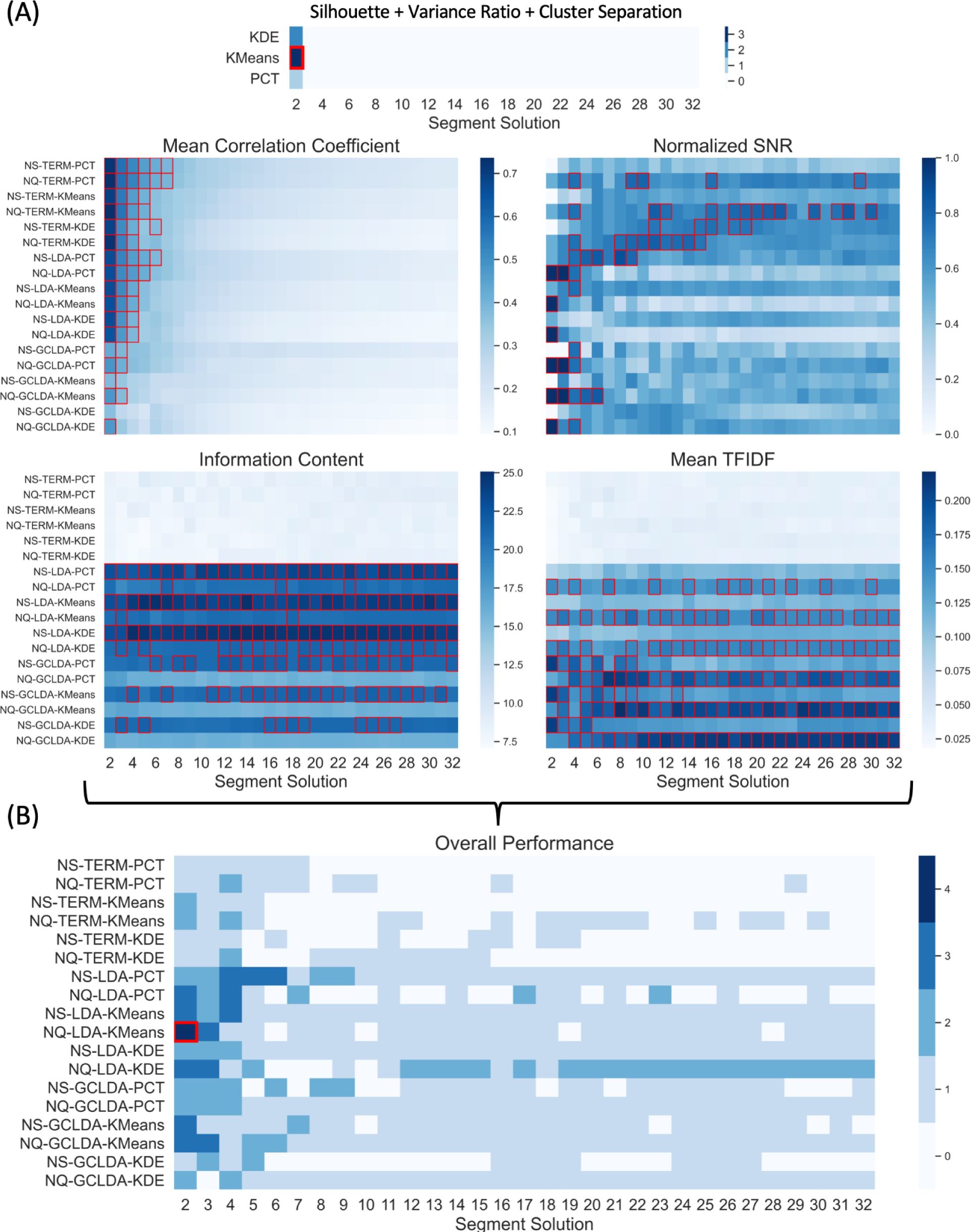
Overall Performance. A) The five heatmaps present the performance of the 18 methods across the 31 segment solutions using three clustering performance metrics and the four benchmark metrics to evaluate the segmentations, decoding algorithms, and meta-analytic database. Strategies that performed over the threshold were marked with a red square using a 90^th^ percentile for the correlation coefficient and SNR and a 70^th^ percentile for IC and TFIDF, which provided a unique solution. B). Summarize the performance of all benchmark metrics. The heatmap was determined by summing the five thresholded binary matrices from the individual heatmaps. The red square marks the unique solution and best-performing strategy (NQ-LDA-KMeans for two-segment solution).

Next, we aimed to generate a figure that delineates the results in a way that most effectively facilitates the interpretation of the functional gradients. **Figure 10** presents the functional decoding of the connectivity gradient for the best-performing strategy (NQ-LDA-KMeans) according to **Fig 8b**. “Non-functional” terms and terms that were associated with a meta-analytic map that did not have a significant correlation (i.e., FDR-corrected p-value > 0.05) with the gradient map were removed from the radar and word cloud plots. The first column shows the principal gradient as the target for decoding (**Fig. 10a**). The second column presents the gradient maps that resulted from the segmentation of the gradient axis (**Fig. 10b**). The last two columns illustrate the result of the decoding algorithm for each gradient map (**Fig. 10c**). The radar plots show the top meta-analytic terms and their corresponding correlation values. The word clouds, on the other hand, provide a more inclusive representation of the terms more likely associated with the corresponding gradient maps. Overall, the proposed figure displays everything in a hierarchical view that helps to describe the gradient from top to bottom, capturing the continuous transition from unimodal primary sensorimotor regions associated with “motor” and “movement” to transmodal regions associated with the default mode network and representative of abstract functioning, as it was first described by Margulies et al. (Margulies et al., 2016).

**Figure 9.**
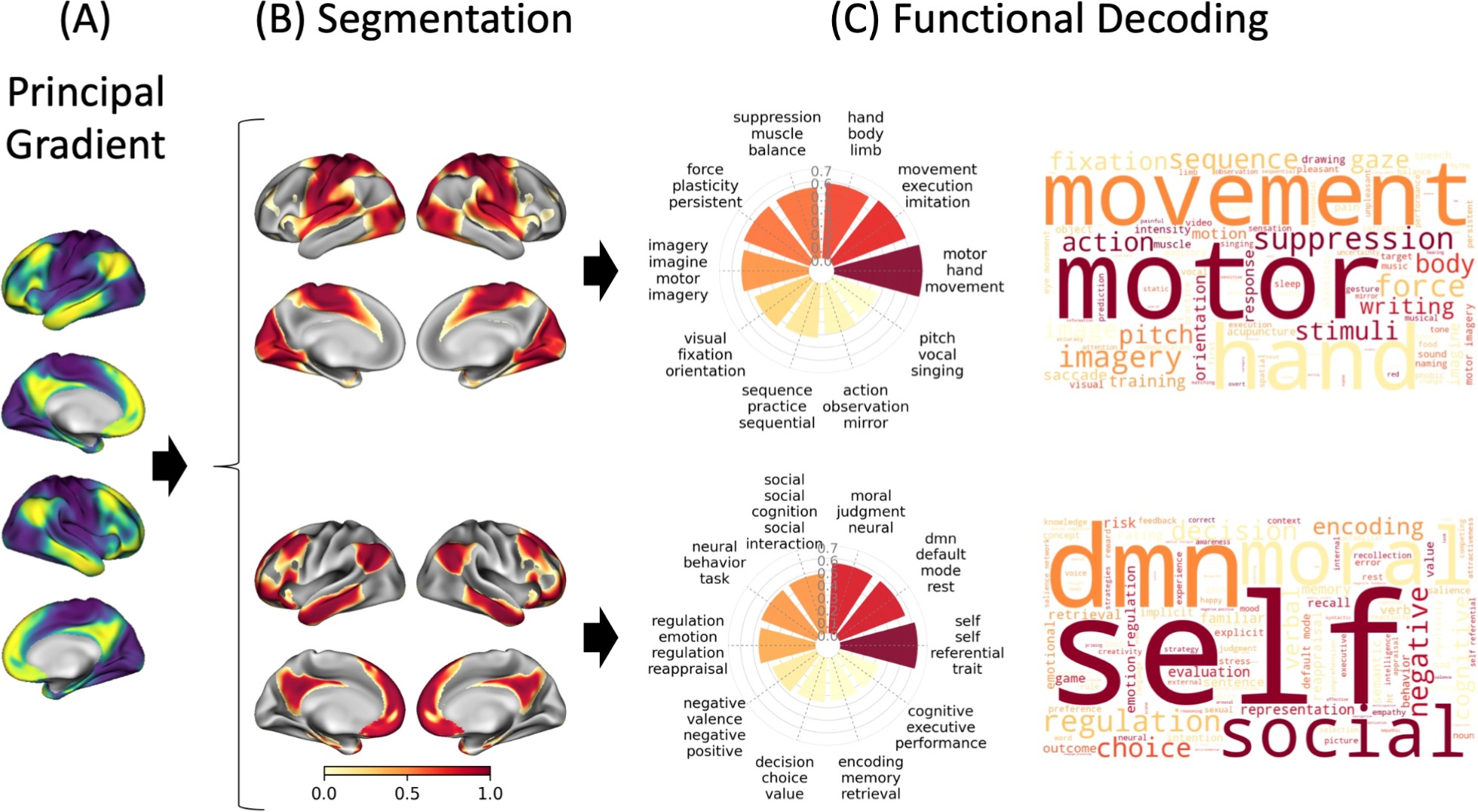
Visualization Approach for Decoding Results of the Principle Gradient of Connectivity. A) Principal gradient of functional connectivity of the cortical surface. B) Segmentation of the principal gradient using a KMeans approach for a segment solution equal to two. C) Results of the functional decoding as shown by a radar plot and a word cloud using an LDA-based meta-analytic map generated using NeuroQuery. Radar plots present the top ten functional terms from statistically significant meta-analytic maps associated with each segment, sorted according to their values, where the values represent correlation coefficients ranging from -1 to 1, with the maps that yielded the highest correlation plotted in the angle equal zero degrees in polar coordinates. Word cloud plots were generated using a frequency estimated by the normalized probability of the topic given the word weighted by the correlation coefficient of the corresponding maps.

**Figure 10.**
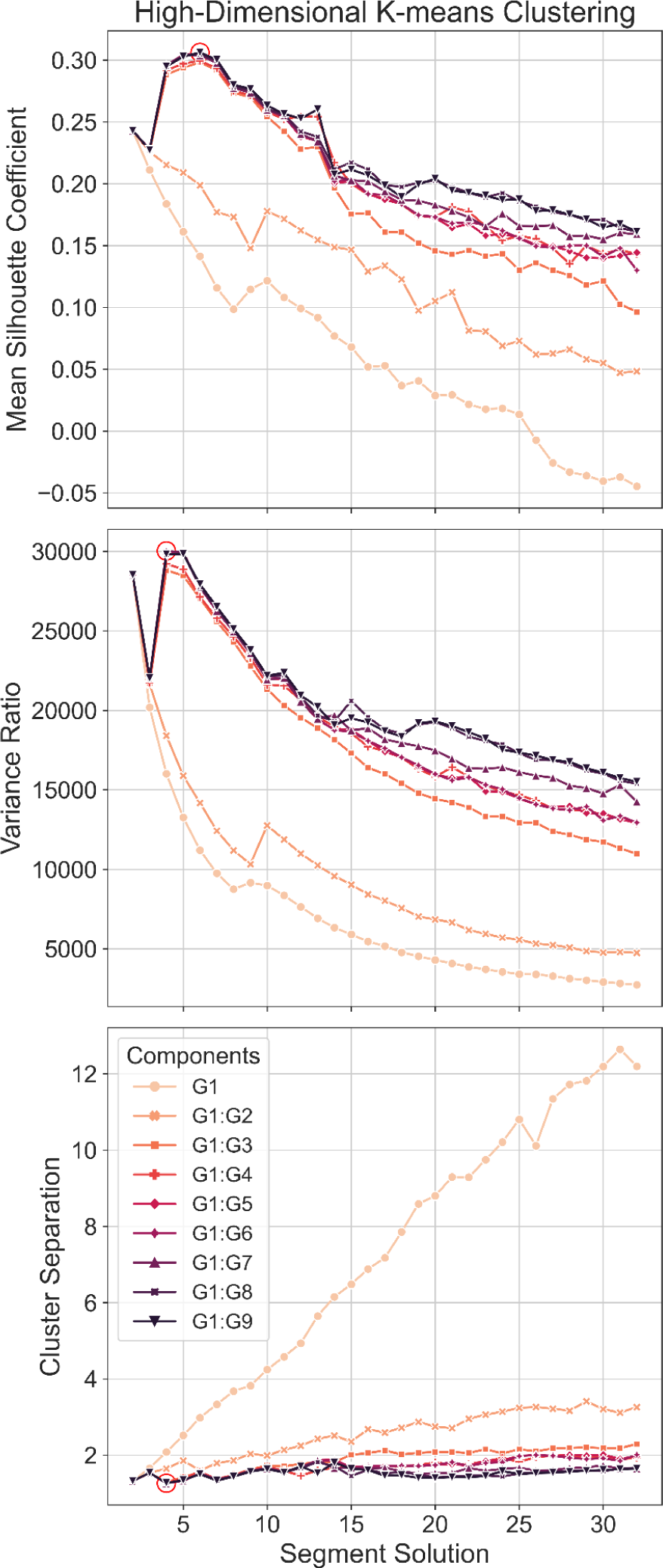
Multidimensional Clustering. Mean silhouette, variance ratio, and cluster separation scores are plotted by segment/cluster solution across gradient dimensions to determine relative performance, with the highest mean silhouette score, highest variance ratio, and lowest cluster separation representing better relative performance (red circle). The segmentation of G1:Gn was performed in the n-dimensional space, i.e., it included gradient 1, gradient 2, …, and gradient n.

### Analysis Step 5: Decoding Multidimensional Gradients

#### Multidimensional Segmentation

Finally, we propose a method for decoding lower-order gradient maps (e.g., second and third gradients), in which all the components are analyzed together in a high dimensional space. To test this method, first, we performed k-means clustering of the gradient for nine different dimensions, ranging from 1D (i.e., only includes the principal gradient: G1) to 9D (i.e., includes the nine first gradients: G1:G9). Mean silhouette scores, variance ratio, and cluster separation were computed to determine the relative performance of the k-means clustering solution across the different dimensions and clustering solutions. The highest mean silhouette score, highest variance ratio, and lowest cluster separation represented the best performance. As illustrated in **Fig. 10**, the best performance was observed when including more than two gradients (2D) in the clustering algorithm, with the best clustering solutions ranging from two to seven clusters. **Fig. S11** summarizes the performance using heatmaps. In particular, we found that a four-cluster solution produced the best solution when including the first seven gradients (**Fig. S11b**). For the next step, as an illustration of high-dimensional decoding, we limited the analysis to 4D space (i.e., the first four components), which has been the focus of attention in the community in recent years (Hong et al., 2020, 2019; Margulies et al., 2016; Paquola et al., 2019). Although the clustering at 7D space performed better, we showed that the four clusters in 4D were very similar (NMI = 0.89) (**Fig. S12**). Clustering similarity was determined using the normalized mutual information (NMI) (Peraza et al., 2020).

#### Multidimensional Decoding

Next, we performed meta-analytic decoding of the four clusters determined by k-means on 4D gradient space. The first four gradients accounted for 75% of the cumulative explained variance. **Fig. 11** presents the result of the decoder in this high-dimensional space. For visualization, we projected the four gradients to 2D space defined by gradients 1 and 3. Additional projections to 2D space combining different gradients can be found in the Supplementary Information (**Fig. S13**). We found cluster 1 to be associated with the default mode network (DMN), cluster 2 with the frontoparietal network, cluster 3 with the visual network, and cluster 4 with the sensorimotor network. Notably, the functional organization defined in high dimensional space was not exactly reproduced by analyzing lower-order gradients separately (**Fig. S14**).

**Figure 11.**
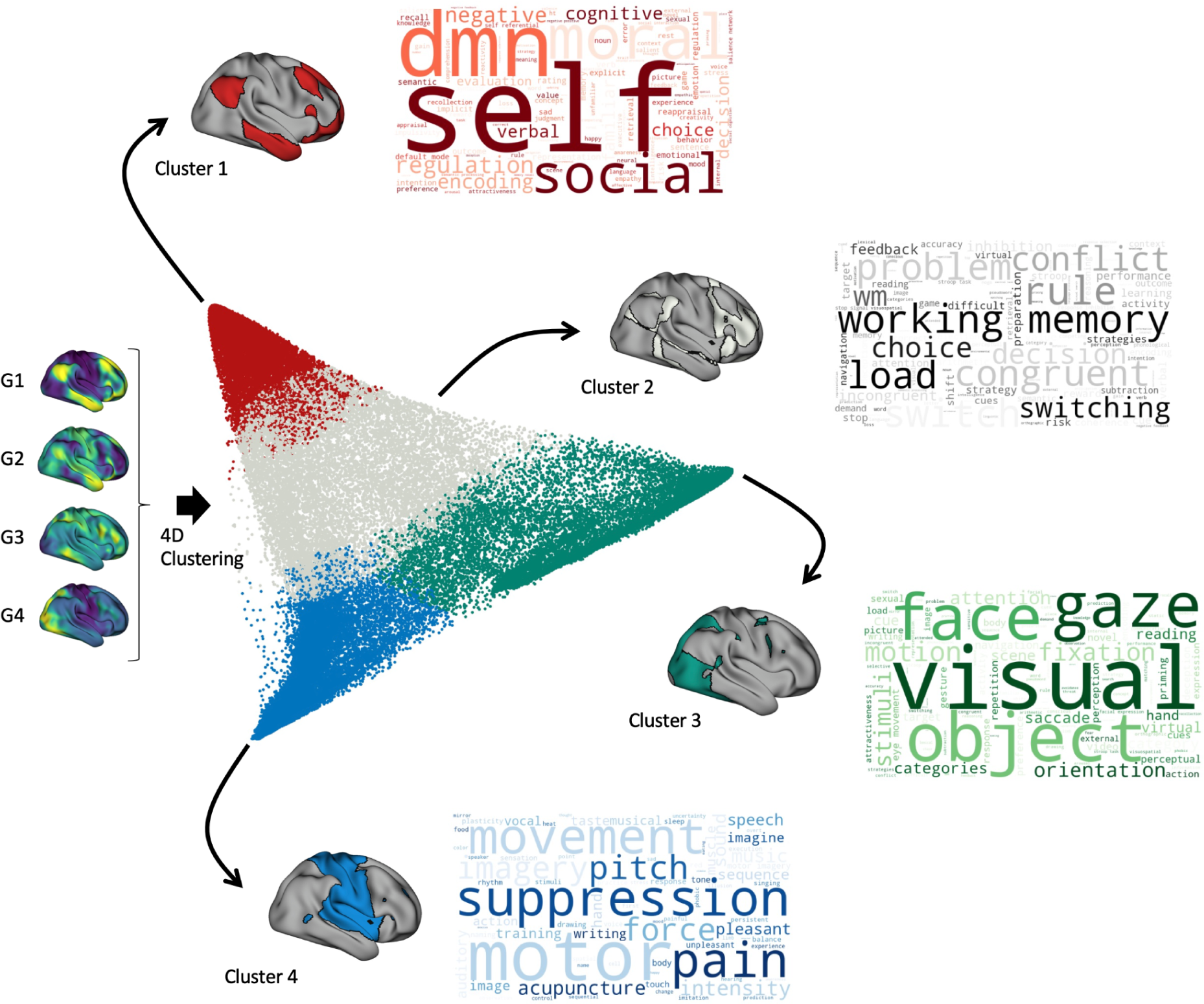
Multidimensional Decoding. Four cluster solutions were determined with the first seven gradients. For visualization, the scatter plot was represented as a projection of the 4D space to the 2D space, determined by gradients 1 and 3. The four clusters are represented in the scatter plot with four distinctive colors. Cluster 1 (red) DMN. Cluster 2 (grays) frontoparietal network. Cluster 3 (green) visual network. Cluster 4 (blue) sensorimotor network. Word cloud plots were generated using a frequency estimated by the normalized probability of the topic given the word weighted by the correlation coefficient of the corresponding maps.

## Discussion

In recent years, investigating macroscale gradients of functional connectivity has become one of the most popular methods for understanding functional brain organization. In the present study, we investigated and sought to improve the framework of data-driven methods for decoding the principal gradient of functional connectivity, thereby promoting best practices for understanding its underlying mechanisms. We evaluated 18 different decoding strategies, which consisted of three segmentation approaches combined with three meta-analytic methods and two databases. First, we found that a data-driven segmentation determined by a k-means algorithm produced the most balanced distributions of boundaries with the highest vertex-wise and mean silhouette coefficient across segment solutions. We also determined that a small number of segments (e.g., two-segment solution) and larger maps (i.e., more than 15% of the brain cortical surface) are preferred for producing maps with high silhouette scores and obtaining high precision in correlation decoders. Furthermore, we observed that an LDA-based decoder, besides showing a high average of top correlation scores, an LDA-based decoder yielded the highest information content, TFIDF, and SNR. Finally, although we noticed minimal differences between the two meta-analytic databases tested in this work, the NeuroQuery database showed a higher SNR, provided by the richness of its vocabulary compared to Neurosynth. The following sections discuss the implications of using a specific segmentation algorithm, meta-analytic decoder, and database for decoding connectivity gradients.

### Segmentation of Connectivity Gradients

The segmentation of the connectivity gradient, the first step in the decoding framework, allows us to characterize the entire gradient spectrum by individually decoding the smaller segments as if they were independent brain maps. This step helps address limitations of standard meta-analytic decoding, which otherwise would yield associations to only vertices at the positive end of the gradient axis, leaving negative coordinates at the other end unexplained. In this study, we demonstrated that the number of segments and the segmentation approach affect the final results of decoding strategies.

First, we found that the segment solution had a major influence on the final result. Indeed, for small numbers of segments, a KMeans algorithm yields the most confident distribution of boundaries, as shown by the silhouette coefficients, variance ratio, and cluster separation. Furthermore, we determined that a large number of segments was detrimental to the performance of correlation decoders, as the average of the top correlation values decreases exponentially with the increase of the segment solution. Although a large number of segments does not directly affect the final result of the decoder after filtering the word to keep only functional terms, and as shown by the word cloud at both ends of the spectrum, a higher number of segments resulted in additional challenges to condense the decoding results for intermediate segments. More importantly, the decoder results for intermediate segments are less reliable than those on both ends, given the low precision of the correlation decoder with high sparsity maps. As a result, we recommend treating results from these subsegments with caution. Prior work has used an arbitrary number of segments ranging from 10 to 20 (Caciagli et al., 2022; Hong et al., 2019; Margulies et al., 2016; Paquola et al., 2019; Wang et al., 2023). Here, we recommend against using that many segments as there was a decrease in the silhouette score and a reduction in the reliability of correlation decoders. Furthermore, many subsegments complicate the classification of segments in the middle of the spectrum without adding extra benefits to characterizing the gradient as a whole. When analyzing the principal gradient of functional connectivity, we recommend using a two-segment solution, given the high silhouette coefficient, good relative performance for correlation decoders, and the absence of intermediate segments.

For higher-segment solutions, the selection of the segmentation algorithm only produced minor effects on the final results. The word clouds showed similar terms regardless of the segmentation algorithm used. Such similarity arises from the stability in clustering solutions for the two end segments of the gradient, especially for a large-segment solution. That similarity was less clear for a small number of segments, where the segment solution was more distinctive across segmentation approaches. Strategies using KDE performed well in their segment assignment as measured by the silhouette coefficient. However, the computation of the boundaries is computationally expensive, especially for a large number of segments. As such, we recommend against using this algorithm. Previous work has used an arbitrary PCT segmentation approach, which yields a well-balanced segment assignment across solutions given the relatively high silhouette coefficients. Ultimately, we recommend using a KMeans algorithm, which yields one of the two highest silhouette coefficients, variance ratio, and lowest cluster separation for the two-segment solution. Furthermore, the k-means algorithm allows for additional components from the dimensionality reduction, which produces an improved gradient segmentation.

### Meta-Analytic Decoder

Starting from the premise that the labels associated with the meta-analytic maps accurately represent the underlying cognitive constructs, our goal was to identify the meta-analytic maps with the highest correlation value to the target map. Further, we expected such maps to be associated with a meaningful collection of terms and thus comprise high information content, TFIDF, and SNR.

As predicted, term-based decoders performed better than LDA and GC-LDA in terms of their correlation values, which is likely a natural outcome for a model with a large number of available maps in comparison to the fixed number of maps for topic-based decoders (i.e., 200 topics maps). Although the top term-based maps accurately matched specific stimuli, their labels are known to be redundant and ambiguous, given that they only consist of a single term. As a result, we hypothesized that such a model offers a limited characterization of the target maps. Here, we demonstrated the veracity of those previous assumptions by showing how the term-based decoder yielded lower information content and TFIDF. On the other hand, topic-based meta-analytic models generated maps labeled with numerous terms per map, thus addressing the redundancy and ambiguity of the single term from term-based decoders. In particular, LDA-based produced meta-analytic maps that yielded a relatively high correlation value and a collection of terms that naturally improved the information content, TFIDF, and SNR. Noteworthy, correlation decoders showed a decrease in precision when the target maps spanned less than 15% of the brain cortical surface.

Although GC-LDA maps exhibited poor performance in terms of correlation coefficient when used in a correlation decoder framework, we think that the GC-LDA decoder, as implemented in the original paper by Rubin et al. (Rubin et al., 2017), will provide a more accurate characterization of the gradient maps. Therefore, we recommend against using GC-LDA maps in a correlation decoder framework. Although term-based decoders perform very well in terms of their correlation coefficient, we suggest using an LDA-based decoder combined with a large number of topics (e.g., between 100 and 200 topics), as it performed reasonably well in terms of their correlation coefficient and also offered a collection of terms for improved characterization of the target maps. Critically, we advise filtering out terms that do not provide functional information, given that anatomical, clinical, or non-specific labels only add noise to the classification.

In the current study, we chose to report only the top three words for topics to simplify the comparisons, given that those three words accounted for most of the probability of all the words within a topic. However, we note that researchers are encouraged to include additional words from a topic if that will improve the characterization of the target map.

### Meta-Analytic Database

Last but not least, the database used to train the models represents a key element of a meta-analytic decoding strategy, as the accuracy of the coordinate extraction and the heterogeneity of terms, combined with the extent of the corpus, will directly affect the performance of a decoder. As such, we showed that NS and NQ performed similarly regarding their correlation profile. We think the similarity is likely due to the resemblance between their corpora, comprising a similar number of overlapping documents. However, we observed that the vocabulary size may help improve a decoder’s information content and produce more functional terms. Therefore, we recommend using a large database with a large and rich vocabulary like NeuroQuery or a combined database that includes studies from Neurosynth and NeuroQuery.

### Decoding Multidimensional Gradients

Finally, we proposed a method for decoding lower-order gradient maps (e.g., second and third gradients), in which all the components were analyzed together in a high dimensional space. Importantly, we recommend against separately interpreting these lower-order gradients. These gradients still contain important information from the functional organization, accounting for approximately 15% of the information from the original connectivity matrix. We demonstrated that the second gradient separated DMN and the frontoparietal network, aligned with previous work (Smallwood et al., 2021). Similarly, the third gradient separated the sensorimotor and visual networks, also aligned with previous work (Margulies et al., 2016). However, decoding these maps alone failed to reproduce the more precise functional organization established in the high-dimensional space. We conclude that this is primarily because the new segments for decoding were defined with limited information from the original connectivity matrix. Alternatively, we suggest using them with the principal gradient to improve the clustering of functional regions, as combined they all account for almost all the variance explained from the original connectivity matrix.

The results show that performing the clustering algorithm on higher dimensions improves all cluster performance metrics tested. In particular, when including more than the first two gradients, we found better cluster solutions, given the high mean silhouette score, high variance ratio, and low cluster separation. The optimal clustering solution was found for four clusters in 7D space. However, we used the four-cluster solution from 4D space to illustrate the decoding analysis, which showed a high NMI to 7D. Functional decoding of the four clusters revealed the functional information of the first four components provided. Cluster 1 and 4 were associated with the DMN and the sensorimotor network, consistent with the two ends of the principal gradient (Katsumi et al., 2023; Margulies et al., 2016). Cluster 2 was associated with the frontoparietal network, usually found in the third component (Katsumi et al., 2023; Smallwood et al., 2021). Finally, Cluster 3 was associated with the visual cortex, which is usually separated from the sensorimotor network when including the second component (Katsumi et al., 2023; Margulies et al., 2016). Importantly, we found that the similarity in the explained variance of lower-order gradients can yield a different ordering of the components. The second and third gradients were switched compared to previous dimensionality reductions of the HPC S900 dense connectome. This ordering issue, however, can be addressed by performing the analysis in the high dimensional space, as suggested in this work, where the ordering of the gradients is irrelevant.

We recognize that the number of components and clusters to consider in an analysis may depend on the application and the type of connectivity matrix from which the gradients are extracted. In this work, we used a group average thresholded connectivity matrix. However, researchers may use other connectivity matrices, including those at the subject level, from atypical participant groups or different development stages, as well as other matrices with negative or unthresholded values. As a main step, we suggest performing a similar multidimensional segmentation for a set of different dimensions and number of clusters and determining the optimal solution for the particular case using clustering performance metrics such as silhouette, variance ratio, and cluster separation scores.

### Derived Products

The workflows and methods proposed in this work have been implemented in `Gradec` (Peraza et al., 2023), a new open-source Python package developed during the design of this project. `Gradec` includes different modules to perform segmentation/clustering, meta-analytic decoding, and visualization of the decoding results. Notably, the decoding algorithms implemented in the cortical surface are not limited to connectivity gradients. They can also be applied to cortical brain maps from different modalities, such as probability maps, statistical maps, and more complex modalities (e.g., inter-subject variability maps).

### Limitations

Several considerations may limit the present study. First, the cortical gradients were estimated on a group-average functional connectivity matrix. However, researchers may also estimate the gradients at the subject level. We postulate that our results will hold for subject-level gradients, given that most group-level gradients are replicable at the individual level (Katsumi et al., 2023). However, future work is needed to verify this assumption. Second, the group-level functional connectivity matrix from the HPC was thresholded at a value that resulted in 10% of edges. Thus, the result of the decomposition may be subject to a positive bias. Further analyses are needed to explore the segmentation and decoding result of gradients estimated on unthresholded connectivity matrices. Third, we only tested the correlation decoder algorithm. Additional analyses are needed to determine if more advanced decoding strategies, such as the native method implemented in the GC-LDA model or the machine learning algorithm Neural Network on Dictionaries (NNoD) (Menuet et al., 2022), are more appropriate to decode gradient maps. Correlation decoder models are limited by design, and although previous research has shown that the labels assigned to a meta-analytic map accurately reflect its cognitive state (Poldrack et al., 2012; Rubin et al., 2017; Yarkoni et al., 2011), we may identify cases where the label does not correctly describe such maps. This result may be found more frequently among non-specific, clinical, or anatomical labels. Although the maps associated with these labels can be filtered out, leaving only functional meta-analytic maps, we cannot be sure all functional maps are correctly assigned.

### Conclusions

In conclusion, we provide recommendations on best practices for gradient-based functional decoding of fMRI data. We found that a two-segment solution determined by a K-means segmentation approach and an LDA-based meta-analysis combined with the NeuroQuery database was the optimal combination of methods for decoding the principal gradient of functional connectivity. We proposed a method for decoding lower-order gradient maps combined with the principal gradient in a high-dimensional space. This combination of approaches and our recommended visualization method for reporting meta-analytic decoding findings will enhance the overall interpretability of macroscale gradients in the fMRI community.

## Acknowledgments

We thank our three anonymous reviewers for their helpful comments and suggestions during the revision of this manuscript. Special thanks to the FIU Instructional & Research Computing Center (IRCC, http://ircc.fiu.edu) for providing the HPC and computing resources that contributed to the research results reported in this paper.

## Ethical Statement

The Human Connectome Project provided the ethics and consent needed for the study and dissemination of HCP data. This secondary data analysis was approved by the Institutional Review Board of Florida International University.

## Data Accessibility

The functional connectivity data were provided by the Human Connectome Project, WU-Minn Consortium (Principal Investigators: David Van Essen and Kamil Ugurbil; U54-MH091657) funded by the 16 NIH Institutes and Centers that support the NIH Blueprint for Neuroscience Research; and by the McDonnell Center for Systems Neuroscience at Washington University.

The meta-analytic databases used in this project are publicly available to download on GitHub. Neurosynth version 7 data release can be accessed at github.com/neurosynth/neurosynth-data; the NeuroQuery version 1 data release can be accessed at (github.com/neuroquery/neuroquery_data)

## Code Availability

This project relied on multiple open-source Python packages, including: BrainSpace (Vos de Wael et al., 2020), Jupyter (Kluyver et al., 2016), mapalign (github.com/satra/mapalign), Matplotlib (Hunter, 2007), Netneurotools (github.com/netneurolab /netneurotools), Neuromaps (Markello et al., 2022), NiBabel (Brett et al., 2020), Nilearn (Abraham et al., 2014), NiMARE (Salo et al., 2022b; RRID:SCR_017398), NumPy (van der Walt et al., 2011), Pandas (McKinney, 2010), PtitPrince (github.com/pog87/PtitPrince), PySurfer (Waskom et al., 2020), RainCloudPlots (Allen et al., 2021), Scikit-learn (Pedregosa et al., 2011), SciPy (Virtanen et al., 2020), Seaborn (Waskom et al., 2017), SurfPlot (Gale et al., 2021), and word_cloud (Mueller et al., 2018). We also used the HCP software Connectome Workbench (`wb_command` version 1.5.0, (Marcus et al., 2011)).

All code required to reproduce the analyses and figures in this paper is available on GitHub at https://github.com/NBCLab/gradient-decoding. High-resolution figures are available through FigShare (https://figshare.com/projects/Meta-analytic_decoding_of_the_cortical_gradient_of_functional_connectivity/172347). The decoding and segmentation modules are available for future re-use here https://github.com/JulioAPeraza/gradec (Peraza et al., 2023), which will be linked to NiMARE (https://nimare.readthedocs.io/en/latest/) as a Python module for gradient meta-analytic decoding in the future. All data and resources that resulted from this paper (e.g., connectivity gradients and trained meta-analytic decoders) are openly disseminated and made available on the Open Science Framework (OSF) at https://osf.io/xzfrt, including the links to the GitHub repository, and figures.

## Authors’ Contributions

ARL, JAP, TS, and MCR conceived and designed the project. ARL, KLB, JEB, JSF, LDHB, RPL, RP, KLR, JLR, and RWL served as annotators for the meta-analytic terms. JAP, MCR, and TS analyzed data. JAP, MCR, TS, KLB, AdlV, JBP, JD, and JDK contributed scripts and pipelines. JAP, ARL wrote the paper, and all authors contributed to the revisions and approved the final version.

