## Supplemental Information for "Methods for decoding cortical gradients of functional connectivity"

### Supplemental Results:

Analysis Step 1: Functional Connectivity Gradient

- Table S1. Characteristics of Decoding Strategies
- Figure S1. Matrix and Gradient of Functional Connectivity
- Figure S2. Cortical Gradients of Functional Connectivity
- Figure S3. Explained Variance Ratio

Analysis Step 2: Segmentation and Gradient Maps

- Figure S4. Segmentation of the Principal Gradient: Boundaries
- Figure S5. Segmentation Similarity
- Figure S6. Overall Clustering Performance
- Segmentation of the Principal Gradient: Gradient Maps
- Silhouette Distribution and Cluster Imbalance
- Confidence Maps

Analysis Step 3: Meta-Analytic Functional Decoding

- Figure S7. Correlation Coefficient Distributions
- Figure S8. Correlation Scores vs Segment Size
- Term-based Meta-Analysis with Neurosynth
- Term-based Meta-Analysis with Neuroquery
- LDA-based Meta-Analysis with Neurosynth
- LDA-based Meta-Analysis with Neuroquery
- GC-LDA-based Meta-Analysis with Neurosynth
- GC-LDA-based Meta-Analysis with Neuroquery
- Uncorrected Correlation Coefficient Distributions
- FDR-Corrected Correlation Coefficient Distributions

Analysis Step 4: Performance of Decoding Strategies

- Figure S9. Proportion of Functional Terms
- Figure S10. Functional Decoder Word Clouds

- Correlation Profiles
- Information Content Profiles
- Term-Frequency Inverse Document Frequency Profiles
- Classifications
- Word Clouds

Analysis Step 5: Decoding Multidimensional Gradients
- Figure S11. Overall High-Dimensional Clustering Performance
- Figure S12. Comparison of Multidimensional Segmentations
- Figure S13. Multidimensional Segmentation
- Figure S14. Meta-Analytic Functional Decoding of Lower-Order Gradients

**Table S1:** Characteristics of Decoding Strategies.

| Method to Generate Meta-Analytic Maps | No. of Maps |
|---|---|
| Term + NS + `CorrelationDecoder` | 3,228 |
| LDA + NS + `CorrelationDecoder` | 200 |
| GC-LDA + NS + `CorrelationDecoder` | 200 |
| Term + NQ + `CorrelationDecoder` | 6,145 |
| LDA + NQ + `CorrelationDecoder` | 200 |
| GCLDA + NQ + `CorrelationDecoder` | 200 |

*Note.* Term, term-based meta-analysis. LDA, Latent Dirichlet allocation-based meta-analysis. GCLDA, Generalized Correspondence LDA-based meta-analysis. NS, Neurosynth. NQ, NeuroQuery.

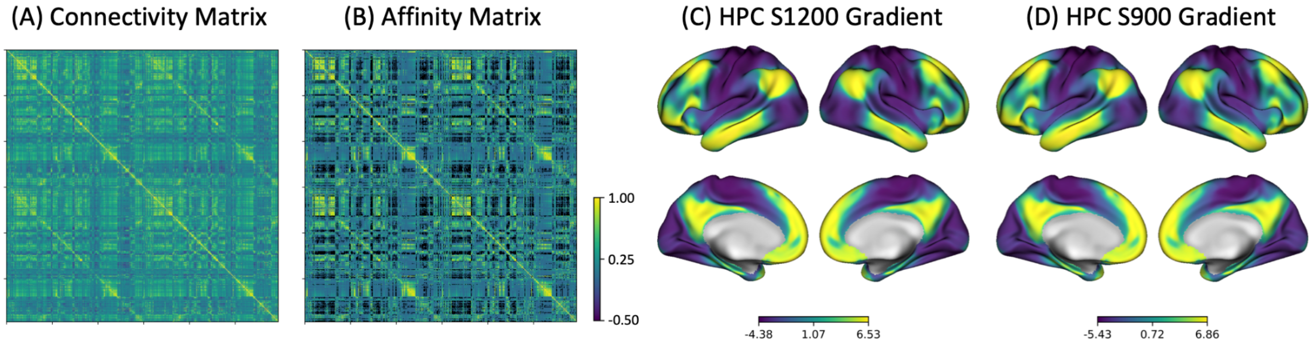

***Figure S1.*** **Matrix and Gradient of Functional Connectivity**. A) Z-scored functional connectivity matrix of the dense connectome HCP S1200 data release. B) Affinity matrix. C) Principal gradient observed in the current paper. D) Principal gradient observed from HCP S900 data release by Margulies et. al. (Margulies et al., 2016).

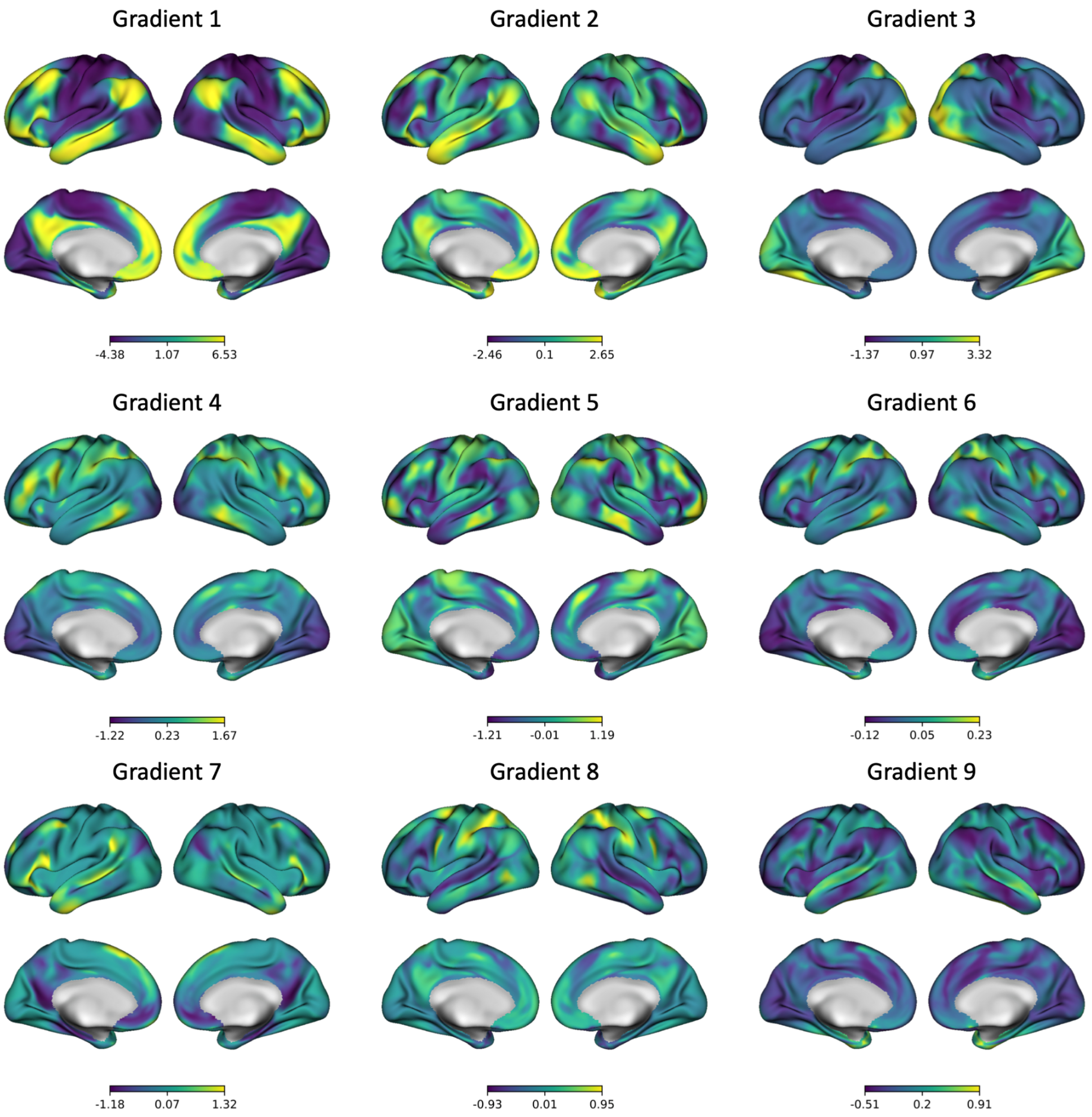

***Figure S2.*** **Cortical Gradients of Functional Connectivity**. The first nine principal components (eigenvectors) from the dimensionality reduction of the functional connectivity matrix.

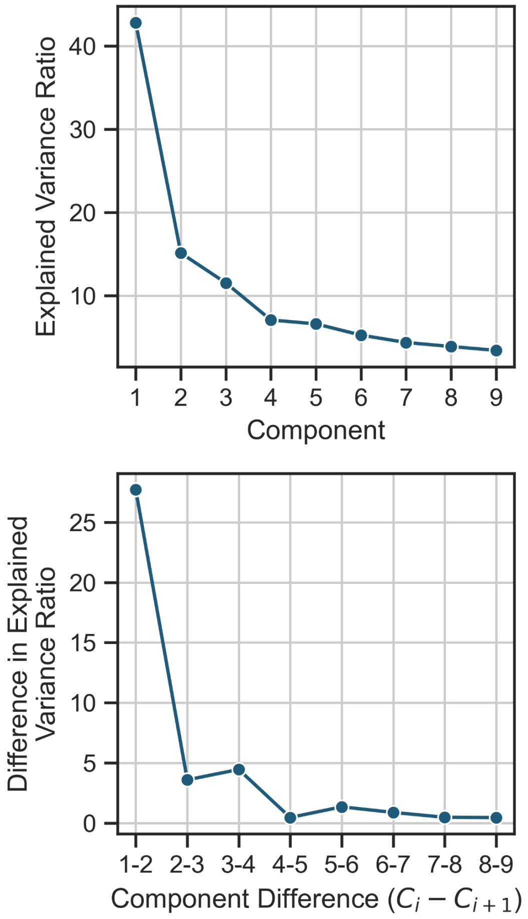

***Figure S3.*** **Explained Variance Ratio**. Top panel: Relative proportion of variance explained (eigenvalues) for the first nine principal components. Bottom panel: Relative proportion of the difference of variance explained for the first nine principal components.

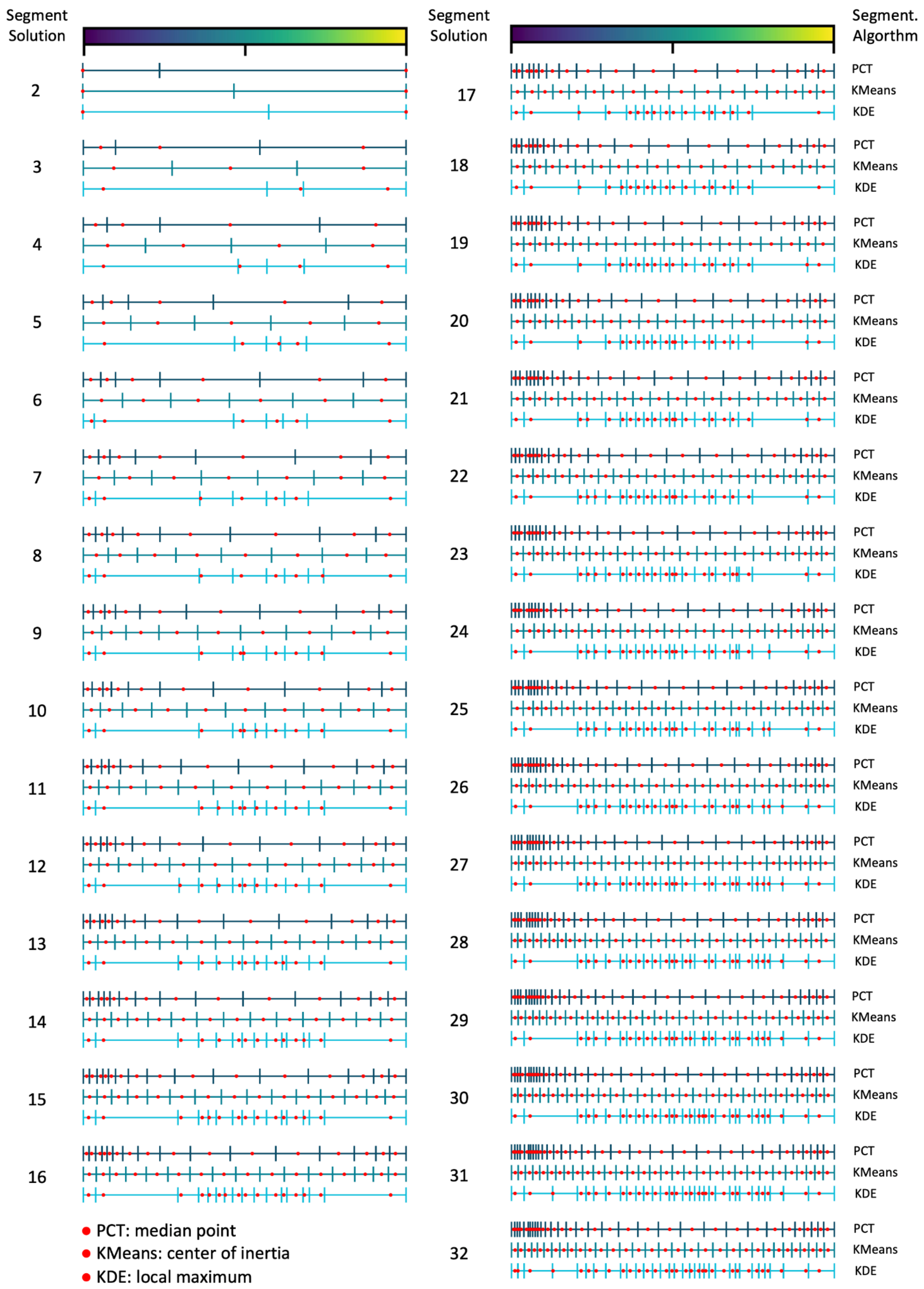

Segment Solution
Segment Solution
Segment. Algorthm
PCT
KMeans
KDE
PCT: median point
KMeans: center of inertia
KDE: local maximum

***Figure S4.*** **Segmentation of the Principal Gradient**. Segmentation of the principal gradient with three different approaches: PCT, KMeans, and KDE. 31 different segmentations were generated, ranging from a solution of two to 32 segments. the red point indicates the peak activation corresponding to the median point, center of inertia, and local maximum for PCT, KMeans, and KDE, respectively

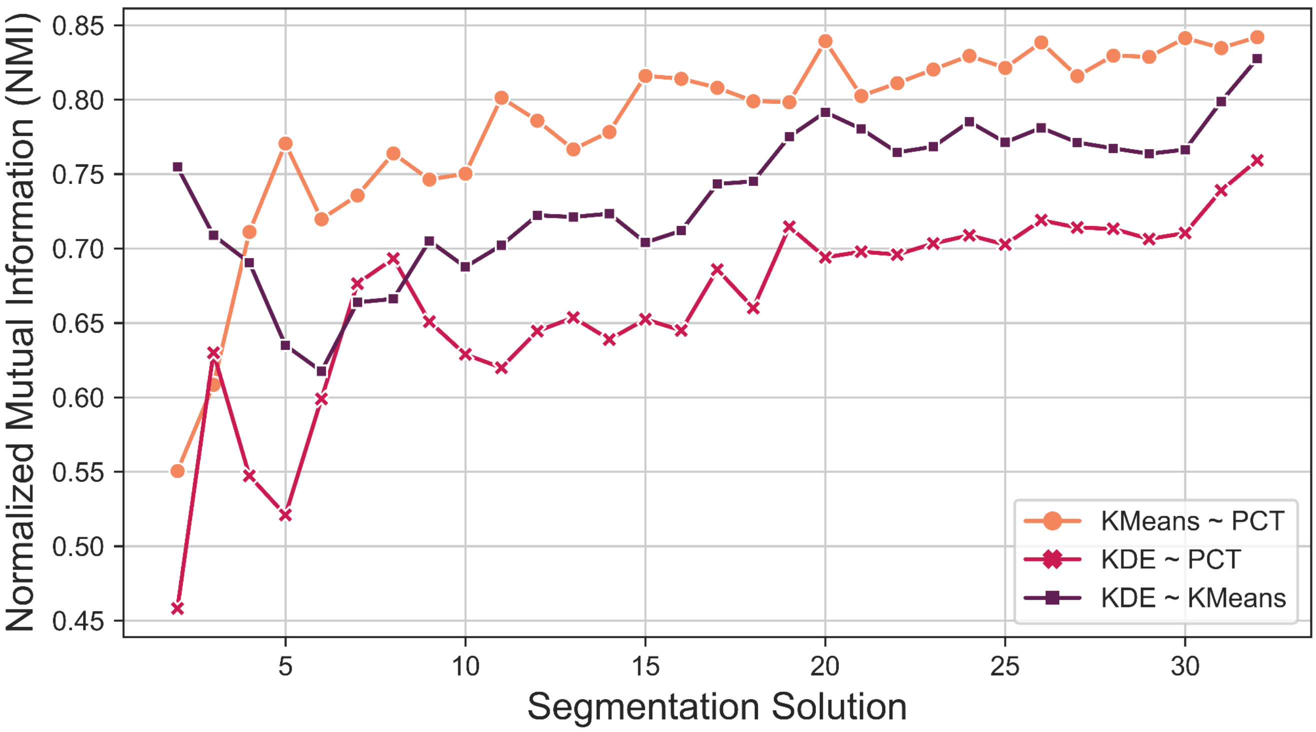

***Figure S5.*** **Segmentation Similarity**. Normalized Mutual Information (NMI) between segmentation approaches across segmentation solutions.

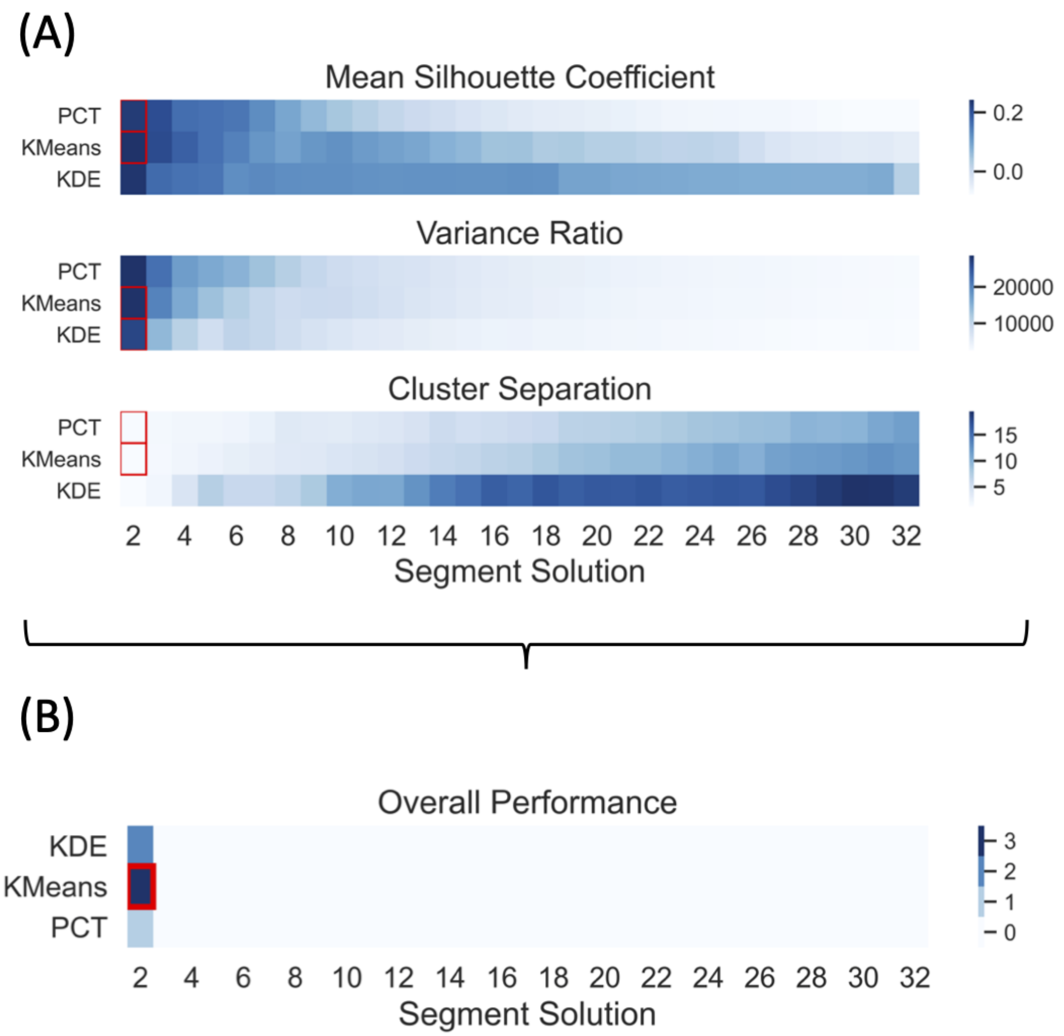

*Figure S6.* **Overall Clustering Performance**. A) The three heatmaps present the performance of the three segmentation approaches across the 31 segment solutions using three clustering performance metrics. Methods that showed performance over the threshold were marked with a red square using a 98$^{th}$ percentile, providing a unique solution to overall performance. B). Summarize the performance of all three benchmark clustering metrics. The heatmap was determined by summing the three thresholded binary matrices from the individual heatmaps. The red square marked the unique solution and best-performing strategy (KMeans for two-segment solution).

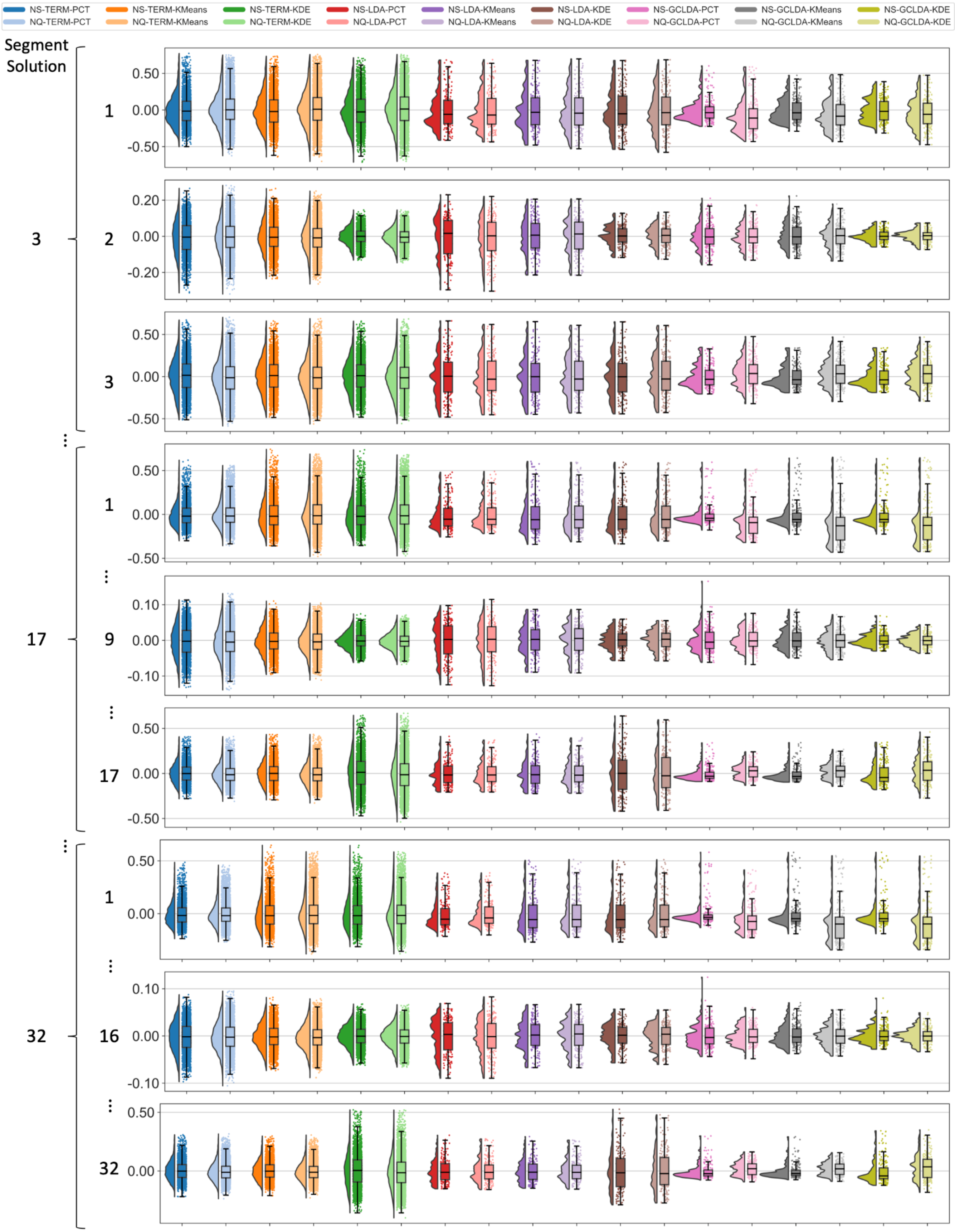
NS-TERM-PCT
NQ-TERM-PCT
NS-TERM-KMeans
NQ-TERM-KMeans
NS-TERM-KDE
NQ-TERM-KDE
NS-LDA-PCT
NQ-LDA-PCT
NS-LDA-KMeans
NQ-LDA-KMeans
NS-LDA-KDE
NQ-LDA-KDE
NS-GCLDA-PCT
NQ-GCLDA-PCT
NS-GCLDA-KMeans
NQ-GCLDA-KMeans
NS-GCLDA-KDE
NQ-GCLDA-KDE
Segment Solution
3
17
32
1
2
3
9
16
0.50
0.00
-0.50
0.20
-0.20
0.10
-0.10

***Figure S7*. Correlation Coefficient Distributions**. CorrelationDecoder estimates across models (i.e., Term, LDA, and GC-LDA), databases (i.e., Neurosynth and Neuroquery), and segmentation approaches (i.e., PCT, KMeans, and KDE) for three distinctive segment solutions (e.g., 3, 17, and 32). For each segment solution, we plotted three subsegments, including the segments at both ends of the spectrum and the segment in the middle. The values in the distribution (y-axis) represent the correlations between meta-analytic maps and gradient maps. Vertex-wise correlation coefficients range from 1 to -1, where positive values indicate high correlation or similar maps, and negative values (in blue) indicate anticorrelations or dissimilar maps. The distribution for the rest of the segment solutions can be found in the Supplementary Results.

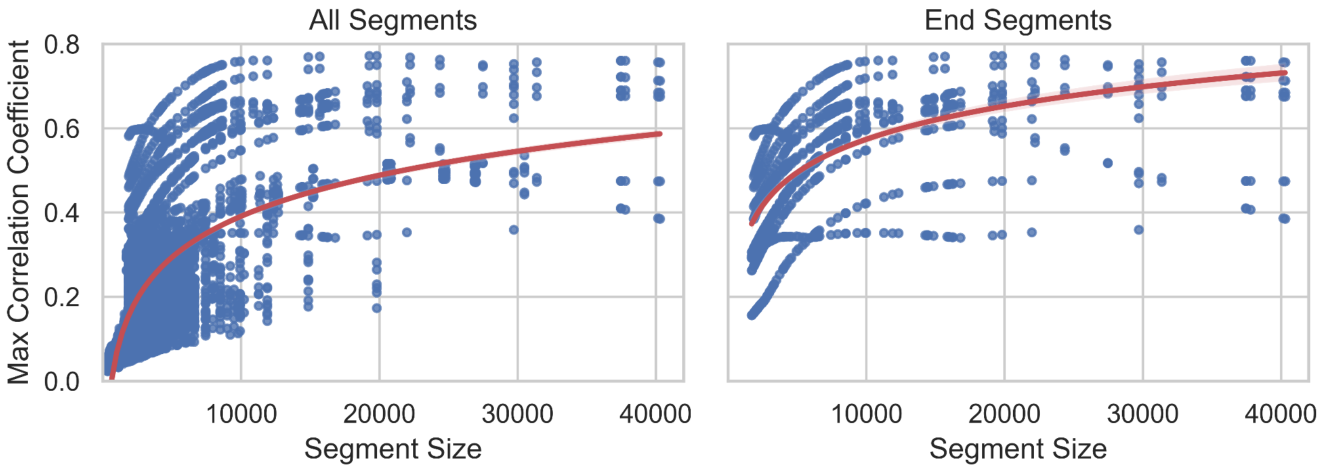

***Figure S8*. Correlation Scores vs Segment Size**. Relation between the correlation coefficients and segment/map sizes. The left panel includes all segments, while the right panel includes only the end segments. The red line shows a fitted logarithmic curve of the data.

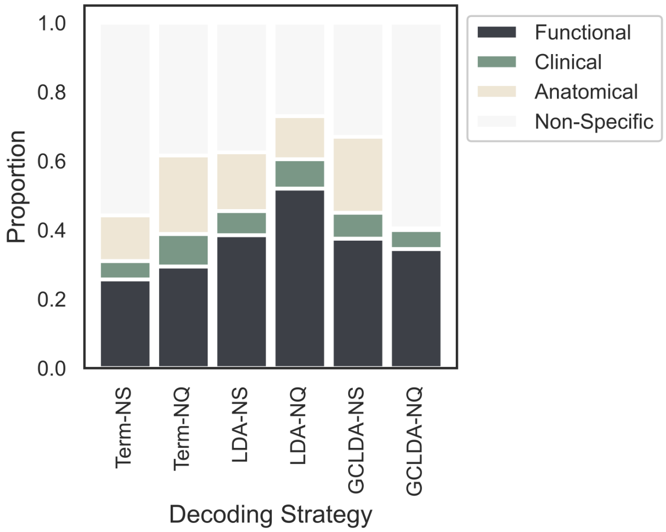

***Figure S9*. Proportion of Functional Terms**. The proportion of "*Functional*", "*Clinical*", "*Anatomical*", and "*Non-specific*" terms/topics are plotted against the 6 different models. Term, term-based meta-analysis. LDA, LDA-based meta-analysis. GC-LDA, GC-LDA-based meta-analysis. NS, Neurosynth. NQ, Neuroquery.

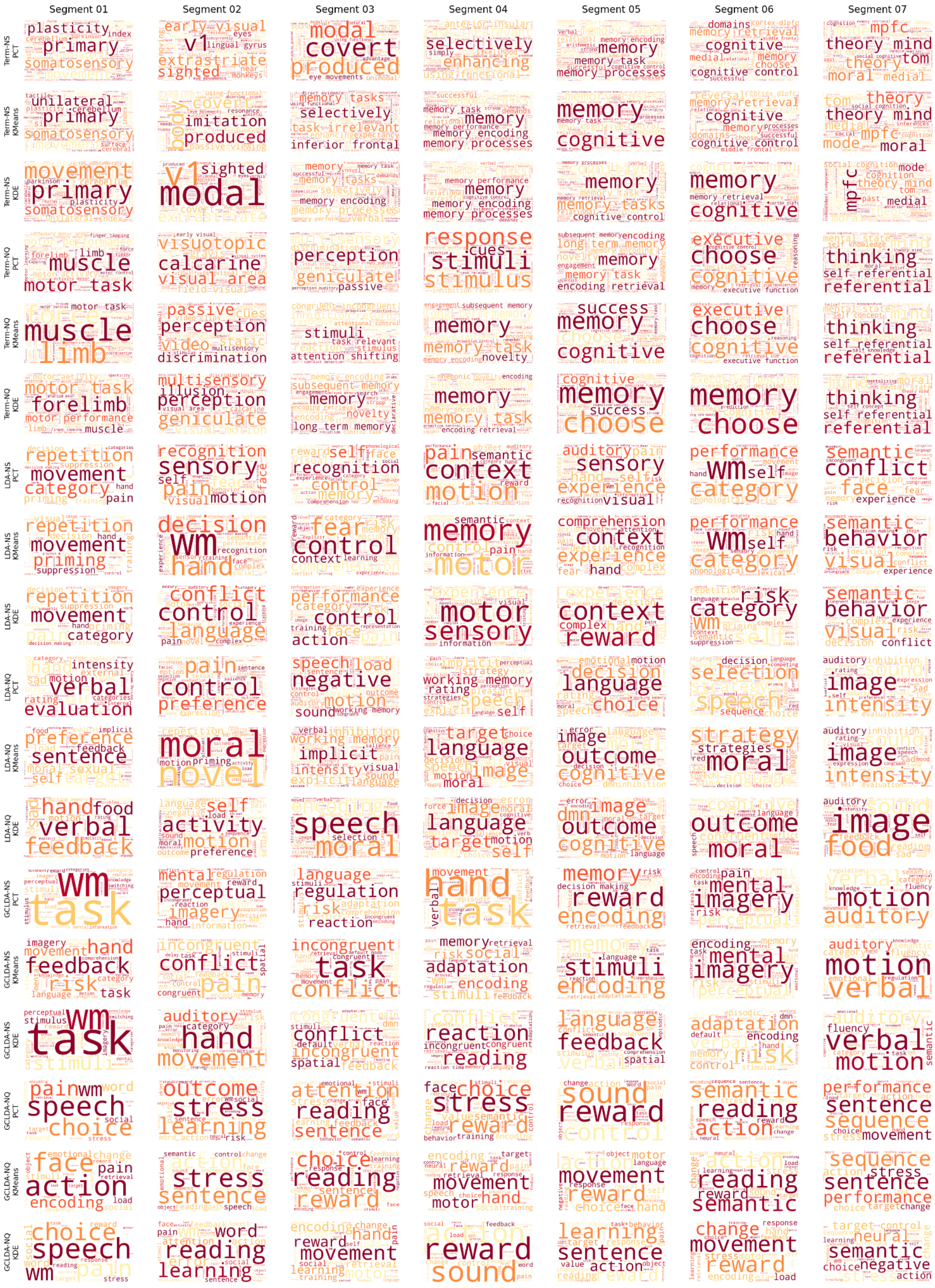

Segment 01
Segment 02
Segment 03
Segment 04
Segment 05
Segment 06
Segment 07
Term-NS PCT
Term-NS KMeans
Term-NS KDE
Term-NQ PCT
Term-NQ KMeans
Term-NQ KDE
LDA-NS PCT
LDA-NS KMeans
LDA-NS KDE
LDA-NQ PCT
LDA-NQ KMeans
LDA-NQ KDE
GCLDA-NS PCT
GCLDA-NS KMeans
GCLDA-NS KDE
GCLDA-NQ PCT
GCLDA-NQ KMeans
GCLDA-NQ KDE

***Figure S10.*** **Functional Decoder Word Clouds**. Word clouds were generated after filtering the results of the top meta-analytic maps and keeping only those classified as "*Functional*". The rows indicate the 18 different strategies for decoding the cortical gradient of functional connectivity that resulted from the combinations of database (NS and NQ), meta-analytic decoder (Term, LDA, and GC-LDA), and segmentation approach (PCT, KMeans, KDE). The columns show the subsegments for a seven-segmentation solution. The frequency associated with each cloud is proportional to the correlation coefficient and the probability of the word given a topic in the case of topic-based decoders. The word clouds for the rest of the segment solutions are linked in the index of this document.

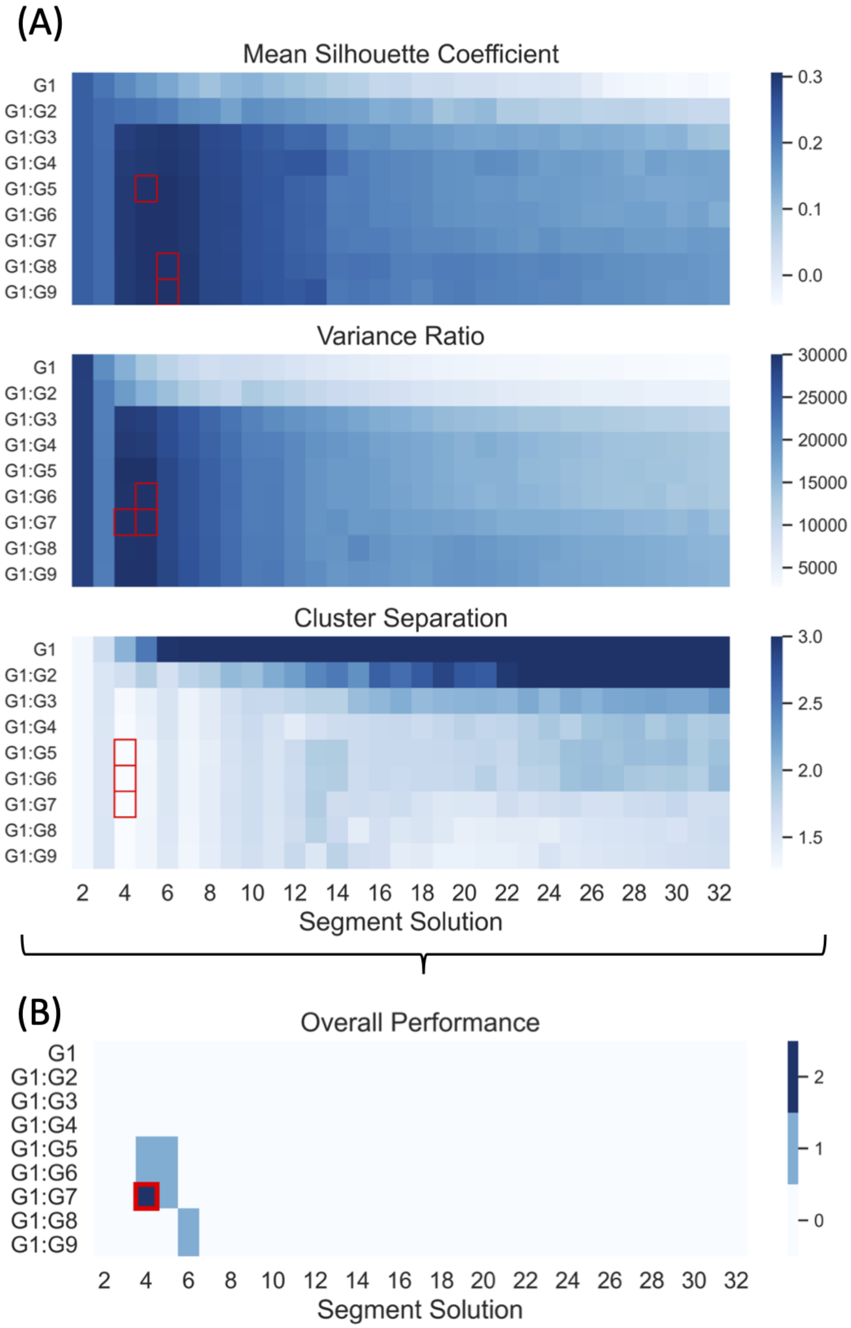

***Figure S11.*** **Overall High-Dimensional Clustering Performance**. A) The three heatmaps present the performance for nine spaces across 31 segment solutions using three clustering performance metrics. Methods that showed performance over the threshold were marked with a red square using a 98 percentile, providing a unique solution to overall performance. B). Summarize the performance of all three benchmark clustering metrics. The heatmap was determined by summing the three thresholded binary matrices from the individual heatmaps. The red square marked the unique solution and best-performing strategy ( four-cluster solution including the first seven gradients of connectivity).

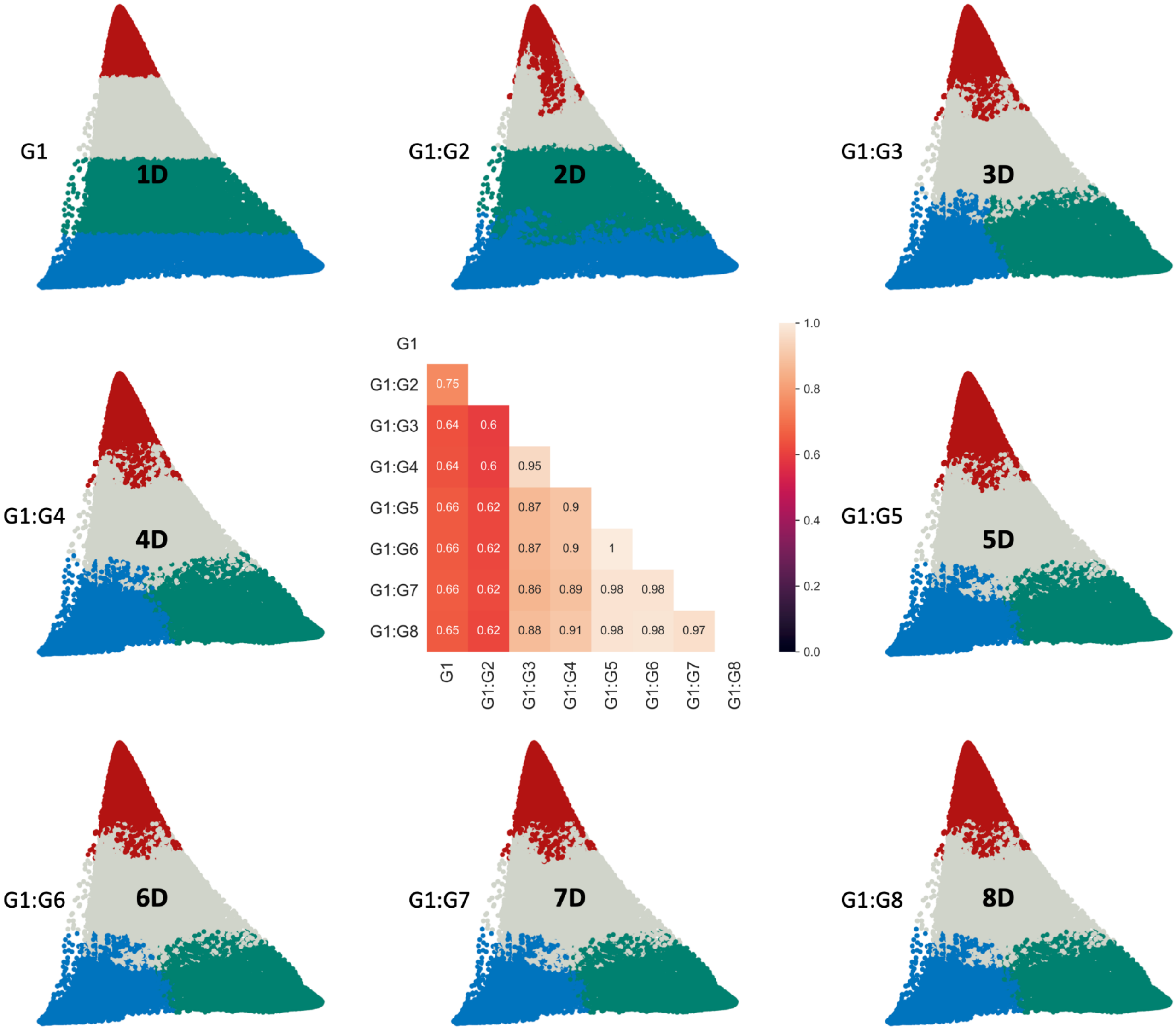

***Figure S12.*** **Comparison of Multidimensional Segmentations.** Four cluster solutions across gradient spaces: from 1D to 8D space. All spaces were projected to a 2D space defined by gradients 1 and 3 to facilitate the visual comparison. The heatmap shows the similarity in the clustering solution as defined by the normalized mutual information (NMI) score.

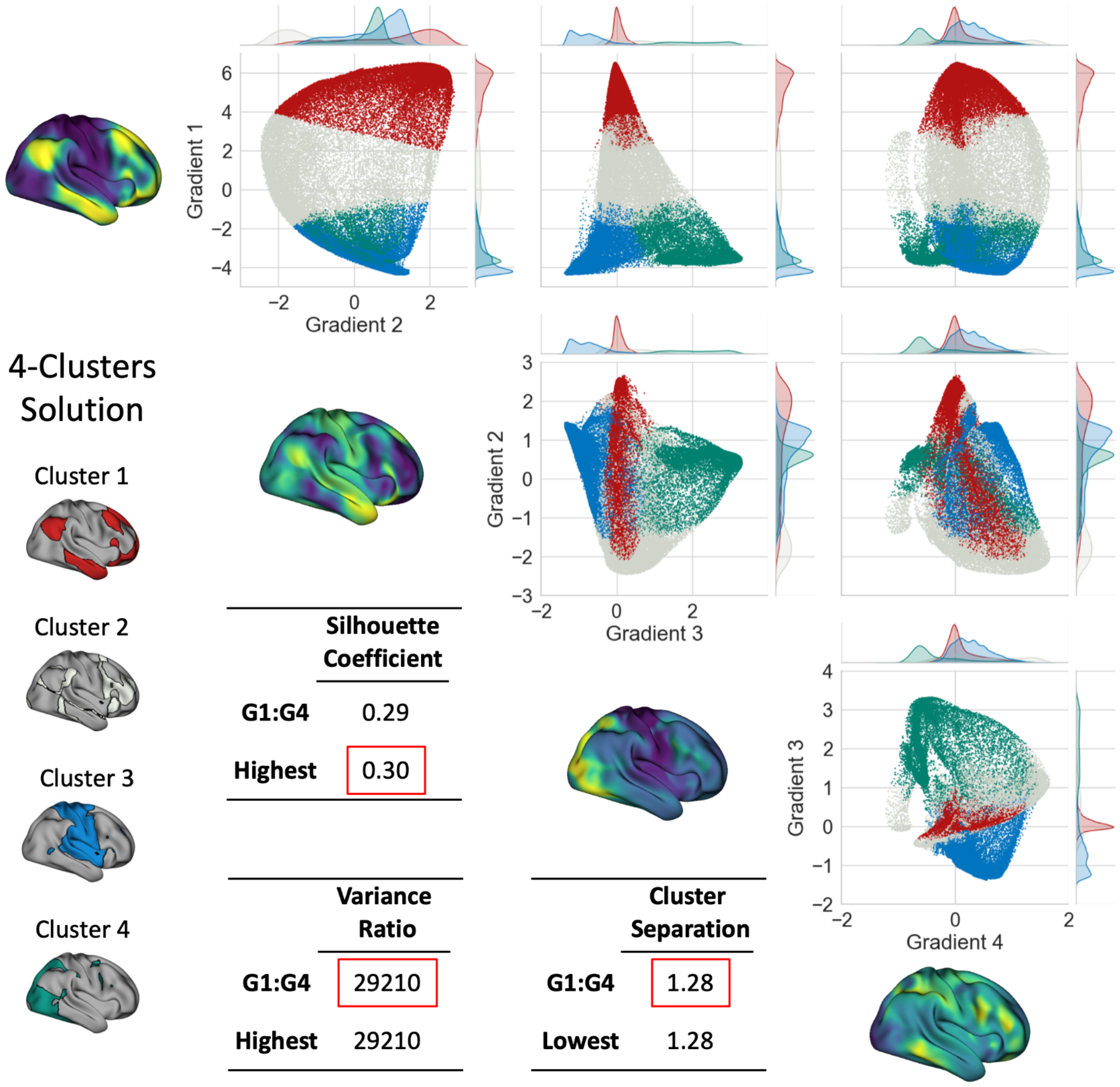

***Figure S13.*** **Multidimensional Segmentation.** Projection of the 4D space to the 2D space defined by the combination of gradients 1, 2, 3, and 4 for visualization. The scatter plots were color-coded based on the four-cluster solution. Cluster 1: red, cluster 2: gray, cluster 3: blue, and cluster 4: green. The three tables present the silhouette, variance ratio, and cluster separation scores. We highlighted the differences in the three scores with the highest score obtained for the four-cluster solution in the 7D space.

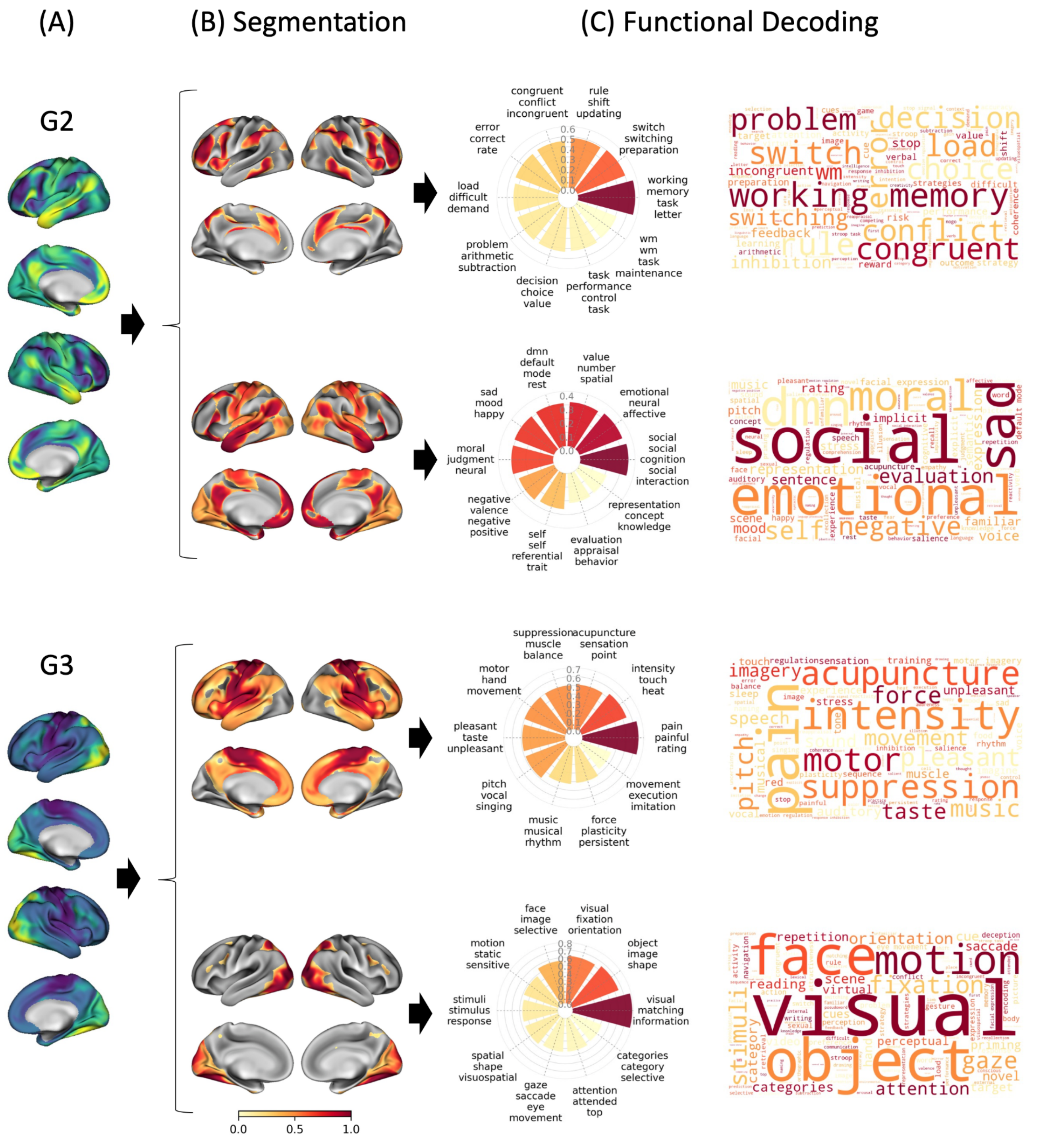

***Figure S14*. Meta-Analytic Functional Decoding of Lower-Order Gradients.** A) Second (G2) and third (G3) gradient of functional connectivity. B) Segmentation of G2 and G3 using a KMeans approach for a two-segment solution. C) Results of the functional decoding using an LDA-based meta-analytic map generated using NeuroQuery. Radar plots present the top ten functional terms associated with each segment, sorted according to their correlation coefficient, ranging from -1 to 1, with the maps that yielded the highest correlation plotted in the angle equal to zero degrees in polar coordinates. Word cloud plots were generated using a frequency estimated by the normalized probability of the topic given the word weighted by the correlation coefficient of the corresponding maps.
